# Repurposing a macromolecular machine: Architecture and evolution of the F7 chemosensory system

**DOI:** 10.1101/653600

**Authors:** Davi R. Ortega, Poorna Subramanian, Petra Mann, Andreas Kjær, Songye Chen, Kylie J. Watts, Sahand Pirbadian, David A. Collins, Romain Kooger, Marina G. Kalyuzhnaya, Simon Ringgaard, Ariane Briegel, Grant J. Jensen

**Author notes:** Rex Richards Building, South Parks Road, Oxford OX1 3QU, UK.

## Abstract

How complex, multi-component macromolecular machines evolved remains poorly understood. Here we reveal the evolutionary origins of the chemosensory machinery that controls flagellar motility in *Escherichia coli*. We first identified ancestral forms still present in *Vibrio cholerae, Pseudomonas aeruginosa, Shewanella oneidensis* and *Methylomicrobium alcaliphilum*, characterizing their structures by electron cryotomography and finding evidence that they function in a stress response pathway. Using bioinformatics, we then traced the evolution of the system through γ-Proteobacteria, pinpointing key evolutionary events that led to the machine now seen in *E. coli.* Our results suggest that two ancient chemosensory systems with different inputs and outputs (F6 and F7) existed contemporaneously, with one (F7) ultimately taking over the inputs and outputs of the other (F6), which was subsequently lost.

## INTRODUCTION

Cells are full of complex, multi-component macromolecular machines with amazingly sophisticated activities. In most cases, how these machines evolved remains mysterious. Presumably they arose through a long series of small steps in which new components and functions accreted onto or replaced old, with fitness advantages at every step that preserved progress along the way. The chemosensory pathway in bacteria and archaea is one such multi-component system. It integrates environmental signals to control cellular functions ranging from flagellum-and pilus-mediated motility to biofilm formation, and chemosensory proteins are key virulence factors for many pathogens. The best-understood function of chemosensory systems is their control of the rotational bias of the flagellar motor, guiding bacteria towards attractants and away from repellents(*1, 2*).

The molecular basis of this activity has been the object of intense study in *Escherichia coli*, where transmembrane <u>M</u>ethyl-Accepting <u>C</u>hemotaxis <u>P</u>roteins, or MCPs, form large arrays at the cell pole(*3*). These chemoreceptors bind attractants or repellents in the periplasm and relay signals to a histidine kinase (CheA) in the cytoplasm(*4*). When activated, CheA first autophosphorylates and then transfers the phosphoryl group to the response regulators CheY and CheB, a methylesterase. Phosphorylated CheY binds to the flagellar motor, changing the direction of flagellar rotation. This allows the cells to switch from swimming forward smoothly (so-called runs) to tumbling randomly. Changes in the duration and frequency of run and tumble phases drive a biased random walk that moves the cells towards favorable environments(*5*). The signal is terminated by a phosphatase, CheZ, that dephosphorylates free CheY(*6*). Phosphorylated CheB tunes the sensitivity of the system by changing the methylation state of the chemoreceptors, opposing the constitutive activity of the methyltransferase CheR(*7, 8*).

While the chemosensory system in *E. coli* is well understood, the structure and function of many others is not. Chemosensory systems have been classified on the basis of evolutionary history into 17 so-called flagellar classes (F1-17), one type IV pili class (TFP) and one class of alternative cellular functions (ACF)(*9*). However, the class names are not reliable predictors of biological role. In *E. coli*, the system that controls the flagellar motor is a member of the F7 class, but in many other bacteria this is not the case. Conversely, in *Rhodospirillum centenum* a member of the F9 class controls biosynthesis of flagella(*10*). Historically, all these pathways have been called chemotaxis pathways in reference to their homology to the biological pathway that gives rise to the chemotaxis phenotype in a diverse set of organisms including *E. coli*. Here we will refer to them instead as chemosensory pathways, to reflect the diversity of outputs that these pathways modulate in response to chemical cues in the environment. 1

In previous work, we and others have used electron cryotomography (cryo-ET) to reveal the *in situ* macromolecular organization of several chemosensory systems(*11–14*). This method allows the study of bacterial cells in a near-native state in three dimensions at macromolecular resolution. Cryo-ET revealed that all the chemosensory systems controlling flagellar motors that have been imaged so far, including the F6 systems of various γ-proteobacteria and the F7 system of *E. coli*, look very similar(*11*). Here, imaging some of these same species under stress, we observed a new kind of chemosensory array. Surprisingly, we identify it as another form of F7, but with a remarkably different architecture compared to that of the canonical *E. coli* F7 system. Tracing its evolutionary history, we find that this novel F7 system actually represents the ancestral form, which in a series of defined steps acquired both the input and output domains of the ancient F6 system to take over control of the flagellar motor, leading to the system seen in modern *E. coli*. The result is a fascinating example of the evolutionary repurposing of complex cellular machinery.

## RESULTS

### *A novel chemosensory array in* V. cholerae *and* P. aeruginosa *in limited growth conditions*

Previously, we used cryo-ET to reveal the structure of two types of chemosensory arrays in *V. cholerae.* When grown in rich medium, the cells contain a polarly-localized membrane-bound array, with a distance of 25 nm between the inner membrane (IM) and the baseplate, composed of kinase and scaffold proteins(*11, 15*). We showed that this array is formed by proteins of the F6 chemosensory system(*16*) (known to control flagellar rotation). In late stationary phase, we found that cells contain another, purely cytoplasmic array consisting of two CheA/CheW baseplates 35 nm apart sandwiching a double layer of chemoreceptors(*16*). This array is formed by proteins of the F9 chemosensory system, but its function is unknown. Here, imaging cells grown into late stationary phase, we observed a third array type. This third type, present in 35% of cells in late stationary phase (Table S1), was membrane-associated and located at the cell pole near the F6 arrays, but was taller than F6 arrays, with a distance between the IM and CheA/CheW baseplate of 38.4±1.9 nm (Fig. 1A).

**Figure 1.**
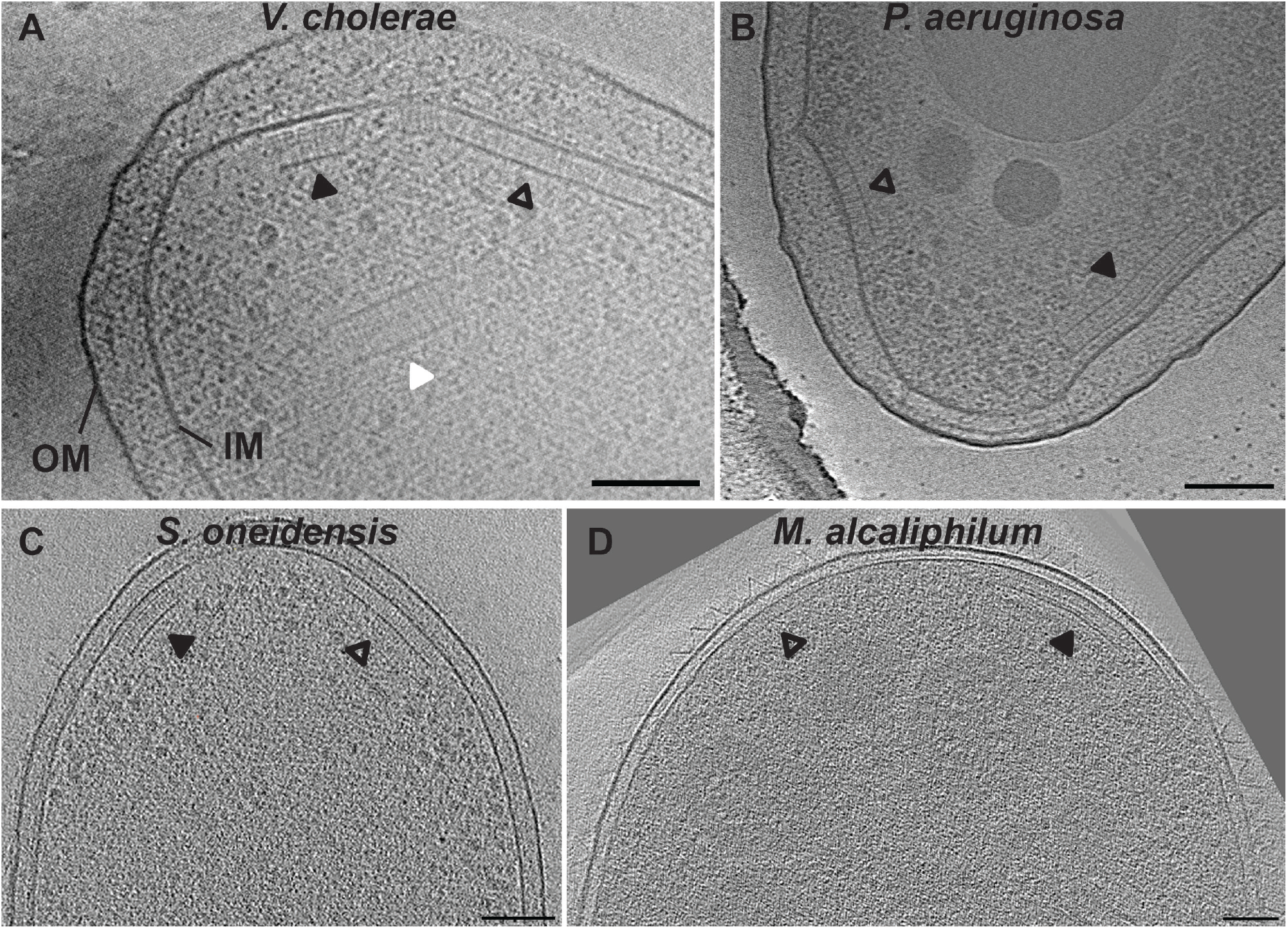
Electron cryotomography reveals that arrays from different putative chemosensory classes, F6 (empty arrows), F7 (black arrows) and F9 (white arrow), have different architectures when observed in side view in various γ-proteobacterial species: (A) *V. cholerae*, (B) *P. aeruginosa*, (C) *S. oneidensis* and (D) *M. alcaliphilum*. Scale bars are 50 nm.

Imaging another γ-proteobacterial species, *Pseudomonas aeruginosa*, grown in nitrogen-limited media, we again observed both short and tall membrane-bound arrays. The short arrays were located at the cell poles, typically in close proximity to the single flagellar motor. The distance between the inner membrane (IM) and the CheA/CheW baseplate was 24.3±1.8 nm. MCPs are classified by length according to the number of heptad repeats they contain(*17*). The length of the shorter array (24 nm) corresponds to receptors belonging to the 40H class that is often associated with F6 systems(*18, 9*), so we assume this array is the F6 system. The additional taller membrane-associated array, present in ~30% of the cells (Table S1), was often (but not always) found at the same cell pole as the putative F6 array and had a distance of 40.3±1.8 nm between the IM and the CheA/CheW baseplate (Fig. 1B).

### The novel array is dependent on proteins of the F7 chemosensory system

To determine which proteins form the novel arrays we observed by cryo-ET, we examined the genomes of *V. cholerae* and *P. aeruginosa*. The genome of *V. cholerae* encodes three chemosensory systems: F6, F7 and F9 (Table S2)(*9*). Having already identified the F6 and F9 systems(*11, 16*), we hypothesized that the novel arrays were formed by the F7 system. The genome of *P. aeruginosa* encodes four chemosensory systems: F6, F7, ACF and TFP (Table S2)(*9*). Since the height of the short array matches that of receptors of the F6 system, and since arrays corresponding to ACF and TFP systems have never been observed in any organism (it is possible that they do not form complexes large enough to be visible in electron cryotomograms), we again hypothesized that the tall array in this organism is formed by the F7 system.

To test this idea, we imaged deletion mutants of F7 genes in *V. cholerae* and *P. aeruginosa* by cryo-ET. Deletion of *cheA* and/or *cheW* of the F7 gene cluster resulted in the absence of the novel tall array (Table S1), confirming that these arrays correspond to the F7 chemosensory system. In both organisms the F7 gene cluster contains two MCPs: one presumably cytosolic class 36H receptor (Aer2/McpB/PA0176 in *P. aeruginosa* and Aer2/VCA1092 in *V. cholerae*), and one receptor of uncategorized class with a predicted transmembrane region (Cttp/McpA/PA0180 in *P. aeruginosa* and VCA1088 in *V. cholerae*). In both species, deletion of the McpA-like receptor had no effect on the presence of the novel array, but deletion of the Aer2 receptor abolished the array (Table S1). We therefore conclude that Aer2 receptors, and not McpA-like receptors, are required for the formation of the F7 arrays.

### F7 chemosensory systems are widespread in γ-Proteobacteria

The histidine kinase CheA gene is used as a proxy to find major chemosensory clusters in genomes(*9*). Using this tool, we selected a non-redundant set of 310 γ-Proteobacteria genomes containing at least one CheA, and found that more than half (176) contained at least one F7 CheA. None of the species we analyzed had more than one F7 system. In the course of other projects, our group has used cryo-ET to image many bacterial species; we found that two of the species in our imaging database(*19*) – *Shewanella oneidensis* MR-1 (*20*) and *Methylomicrobium alcaliphilum* 20Z(*21*) – were γ-Proteobacteria with F7 systems.

The genome of *S. oneidensis* contains two chemosensory systems, one from the F6 class (SO_3200-SO_3209) and another from the F7 class (SO_2117-SO_2126, Table S2) (*9*). In cryotomograms of *S. oneidensis* cells grown anaerobically in batch culture, we observed a single membrane-bound array, usually located at the cell pole in close proximity to the single flagellar motor. The distance between the IM and the CheA/CheW baseplate was 24.5±2.7 nm, as expected for 40H chemoreceptors of the F6 system(*18*). When cells were grown anaerobically in continuous flow bioreactors, however, we observed the novel taller array type in ~10% of cells, often (but not always) at the same cell pole as the F6 array (Table S1). This array had a distance of 35.5±2.7 nm between the IM and the baseplate (Fig. 1C).

Chemotaxis in *M. alcaliphilum* has yet to be explored, but genome analysis predicts three chemosensory gene clusters, one each from the F6 (MEALZ_3148 - MEALZ_3158), F7 (MEALZ_2869-MEALZ_2879) and F8 classes (MEALZ_2939 - MEALZ_2942, Table S2). Cryo-ET of *M. alcaliphilum* revealed two array types: putative F6 arrays with 25.6±2.8 nm between the IM and the CheA/CheW layer, as expected for 40H chemoreceptors(*18*), and, in 25% of cells, taller arrays with a distance of 35.1±2.8 nm between the IM and baseplate (Fig. 1D). The F8 chemosensory system uses a class 34H chemoreceptor with two transmembrane regions. Given the domain architecture in the cytoplasmic portion of the sequence, we expect arrays formed by these receptors to exhibit a distance of ~22nm between the inner membrane and the CheA/ CheW baseplate(*11*). We did not observe any such array in our cryotomograms. We therefore assume that the taller arrays are F7. Given these results in *V. cholerae, P. aeruginosa, S. oneidensis* and *M. alcaliphilum*, we conclude that the tall F7 arrays are wide-spread across γ-Proteobacteria.

### F7 array architectures correspond to the domains of their Aer2-like receptors

In typical F6-like membrane-bound arrays, including all those imaged by cryo-ET in this and previous studies, a layer of periplasmic domains is visible just outside the IM(*11*). In contrast, the novel F7 arrays lacked discernable periplasmic densities. Instead, they exhibited multiple cytoplasmic layers between, and parallel to, the IM and the CheA/CheW baseplate. To better visualize these additional layers, we computed the average 1D profile of electron density from the CheA/CheW layer to the IM in each species (Fig. 2A). Starting from the CheA/CheW layer and moving toward the membrane, *P. aeruginosa, S. oneidensis* and *M. alcaliphilum* showed a density layer very close to the CheA/CheW layer, which we refer to as signaling layer SL. All four species then exhibited two higher layers we name L1 and L2. Finally, all but *M. alcaliphilum* presented an additional layer, L3, near the IM.

**Figure 2.**
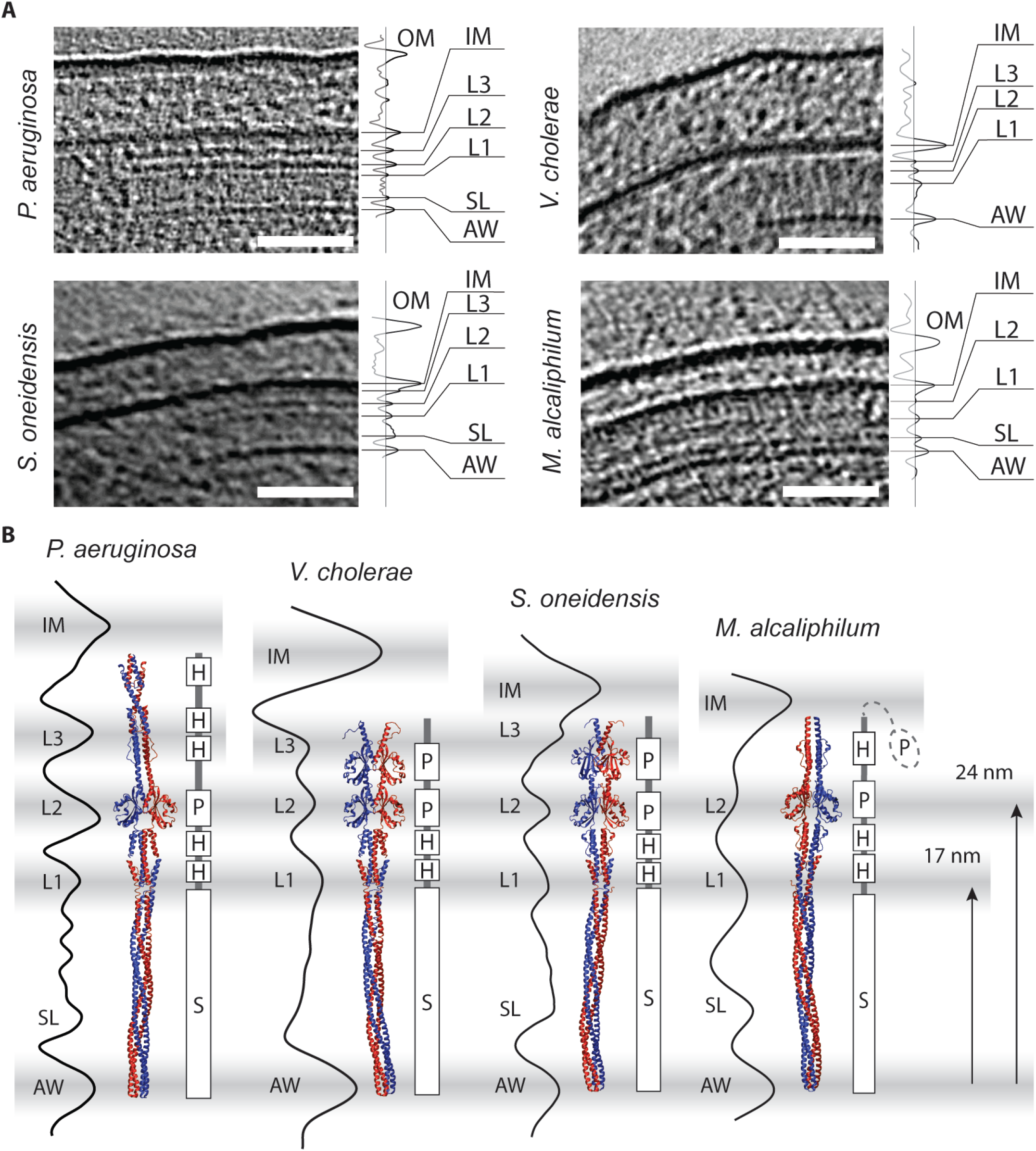
(A) Electron density profiles of chemosensory array side-views reveal intermediate electron density layers in *P. aeruginosa, V. cholerae, S. oneidensis*, and *M. alcaliphilum*. (B) The electron density layers match protein domain features present in homology models of Aer2-like receptor homologs. Note the uncertainty in the localization of the N-terminal PAS domain in *M. alcaliphilum* due to small predicted transmembrane regions (see text).

We wondered whether these density layers we observed by cryo-ET in the novel F7 array corresponded to structural features of its Aer2-like receptor. The *P. aeruginosa* Aer2 receptor is well characterized: it consists of three N-terminal HAMP domains, followed by a PAS domain, two additional HAMP domains and a cytoplasmic signaling domain(*22, 23*). The *V. cholerae* Aer2 receptor is also well characterized: it consists of two N-terminal PAS domains, two HAMP domains and a cytoplasmic signaling domain(*24*). We used CD-VIST(*25*) to predict the domain architecture of the related receptor in the two remaining species. In *S. oneidensis*, the Aer2-like receptor consists of two N-terminal PAS domains, followed by two HAMP domains and a cytoplasmic signaling domain, like Aer2 from *V. cholerae*. As in *P. aeruginosa*, the receptor lacked any discernable transmembrane region and was predicted to be cytosolic. The *M. alcaliphilum* Aer2-like receptor was predicted to contain an N-terminal PAS domain followed by a single HAMP domain, another PAS domain, two HAMP domains and finally a cytoplasmic signaling domain. Unlike in the other species, though, this receptor was predicted to contain two short potential trans-membrane regions (10 and 14 residues) between the N-terminal PAS domain and the rest of the protein. PAS domains are rarely periplasmic(*26*), however, suggesting that even if these regions are embedded in th3e membrane, they do not span it.

Using this information, we constructed a homology model of the Aer2-like receptor in each species based on available atomic models of the protein domains and assuming that the individual domains stack linearly within the four-helix bundle of the receptor dimer. We then manually aligned the homology model for each organism with the corresponding electron density profile (Fig. 2B). In all four cases, the receptor fit well between the CheA/CheW baseplate and IM. We also observed a correlation between densities observed by cryo-ET and domain features of the receptors. In all four species, the first layer, L1, corresponded to the boundary between the cytoplasmic signaling domain and its proximal HAMP domain 17 nm above the CheA/CheW baseplate. The L2 layer 24 nm above the CheA/CheW baseplate corresponded to the PAS domain present in all four species. The second PAS domain present in *V. cholerae* and *S. oneidensis* approximately correlated with the L3 layer 30 nm above the CheA/CheW baseplate. *P. aeruginosa* does not have a second PAS domain; its L3 layer instead appeared to match a HAMP domain (31 nm). The distances from the CheA/CheW baseplate to each layer for each species are listed in Table S3. These correlations further support the conclusion that the novel tall arrays are formed by the Aer2-like MCPs of the F7 system.

### Evolution of the F7 system in Proteobacteria exhibits distinct stages

These findings present a puzzle: the F7 arrays in *P. aeruginosa, V. cholerae, S. oneidensis* and *M. alcaliphilum* all look similar to one another, but very different than the F7 systems we had observed previously in *E. coli* and *S. enterica*. Instead, the *E. coli* and *S. enterica* F7 systems resemble the F6 systems we saw in *P. aeruginosa, V. cholerae, S. oneidensis* and *M. alcaliphilum*. This prompted us to explore the evolutionary relationship between these systems.

To do that, we first constructed a phylogenetic tree of proteobacterial F7 systems using concatenated sequences of CheA, CheB, and CheR (Figs 3 and S1). We built this tree using a non-redundant set of 262 CheABR sequences from F7 (168) and F8 (94) systems from 246 genomes. Sequences from F8 systems were included to root the tree since F8 systems share a common ancestor with F7 systems(*9*). To track the evolution of the system, we grouped monophyletic branches into clades. The first clade comprised all the *ε*-proteobacterial systems, the second all the *α*-proteobacterial systems, the third and fourth non-enteric γ-proteobacterial systems, the fifth and sixth β-proteobacterial systems, and the seventh the enteric γ-Proteobacteria. Thus, remarkably, the CheABR tree was mostly congruent to the general organization of Proteobacteria classes(*27, 28*). This means that the F7 system has been stably associated with its cellular lineage for almost 2.8 billion years(*28*), with either very few horizontal gene transfers or the results of such shuffling events going extinct. To track the distribution of the F7 system through these clades, we built a phylogenetic profile using a new random set of 162 γ-Proteobacteria, in addition to the 4 species imaged in this work and 10 β-Proteobacteria to serve as an outgroup (Fig. S2).

**Figure 3.**
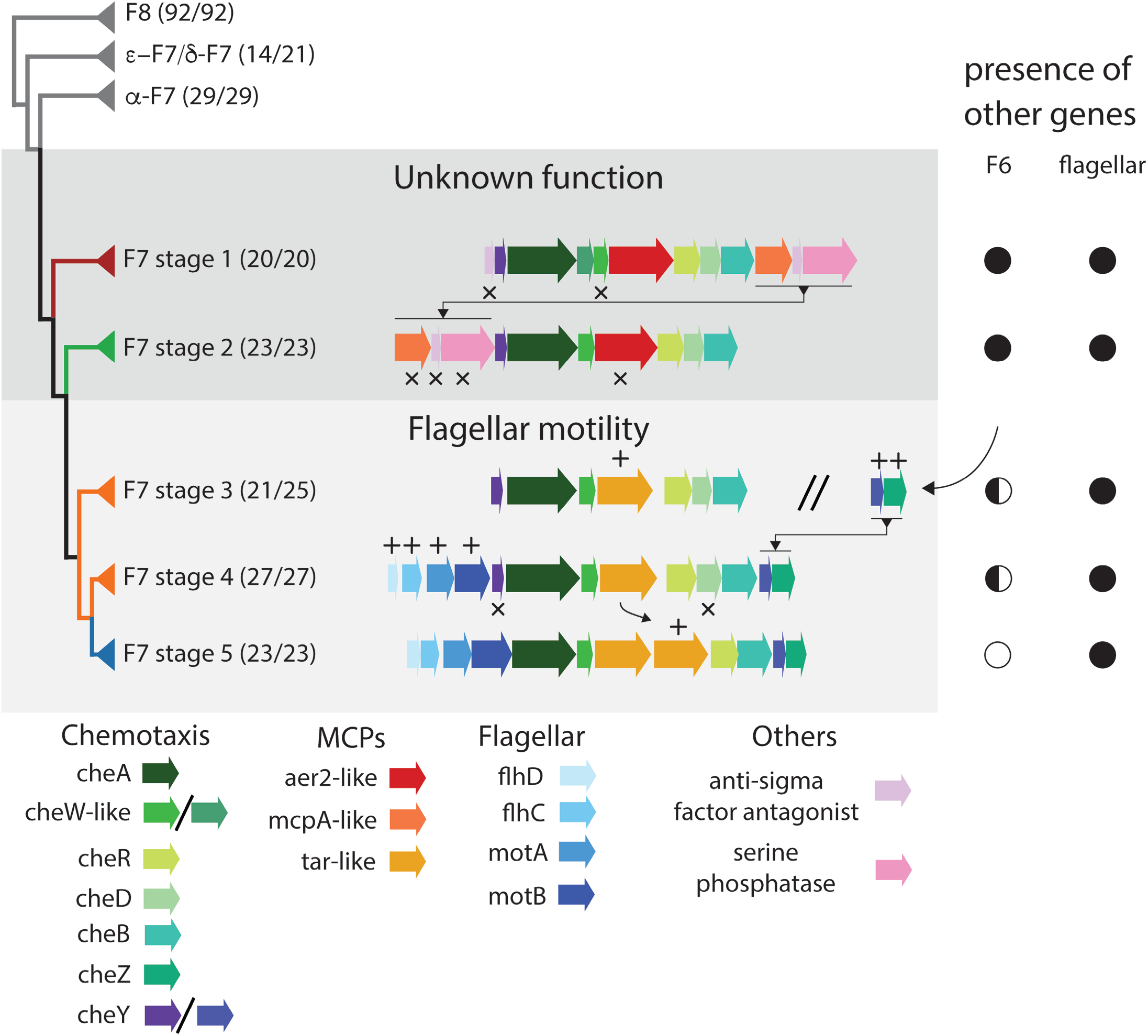
Major events in the evolutionary history of the F7 system in γ-Proteobacteria. Each major branch is identified by the type of its F7 system, and shading indicates the function of the system. Numbers in parentheses indicate the number of genomes from the designated class compared to the number of genomes present in the branch. Differences in these numbers indicate lateral gene transfers and are highlighted in Fig. S1. The “//” symbol is used to show that the protein clusters on each side of the symbol are located in different part of the genome, the symbols plus and crosses indicates genes additions and losses respectively. The presence and absence of other relevant systems in the genomes containing each stage are marked as complete (full circle), partial (half circle) or absent (empty circle).

Next we analyzed the arrangement of genes in the F7 gene cluster in γ-and β-proteobacterial species. We found a specific and characteristic organization of the F7 gene cluster in each of the last five clades of the phylogenetic tree. For clarity, we will refer to these as the first through fifth evolutionary “stages” of the F7 system. The most ancient of these groups, stage 1, was marked by the presence of an anti-sigma factor antagonist followed by *cheY, cheA*, two *cheW*-like genes, an *aer2*-like chemoreceptor, *cheR, cheD, cheB*, an *mcpA*-like chemoreceptor, another anti-sigma factor antagonist and a serine phosphatase. All organisms with a stage 1 F7 system also contained an F6 system elsewhere in their genome (Fig. 3 and Fig. S2). In the transition to stage 2, the most N-terminal anti-sigma factor antagonist of the gene cluster and one of the *cheW*-like genes were lost and the last three downstream genes moved to the front of the chemosensory cluster. Again, all organisms with a stage 2 F7 system had a complete F6 system. Next, in stage 3 these same three (now upstream) genes, including the *mcpA*-like receptor, were lost and an *aer2*-like receptor gene was replaced by a chemoreceptor with two transmembrane regions, a periplasmic sensory domain like those found in well-studied model receptors in *E. coli* (*tar, tap, trg* and *tsr*) and lacking other cytoplasmic protein domains except HAMP and MCP signaling (Fig. S3). In this transition, changes also occurred in the F6 gene cluster: only seven of 21 stage 3 genomes still had an F6 CheA, and none had other core proteins like CheB and CheR, suggesting that the F6 CheA may no longer be functional. All species, however, maintained the F6 *cheY*/*cheZ* pair somewhere in the genome. In stage 4, four flagellar genes (*flhD, flhC, motA* and *motB*) moved to the front of the F7 cluster and the *cheY*/*cheZ* pair previously associated with the F6 system moved to the back. As a result, the F7 cluster now had two *cheY* genes: one from the original F7 system and another from the F6. Further losses occurred in F6 genes: only three of 27 stage 4 genomes retained the F6 CheA (also with no F6 CheB or CheR). Finally, in stage 5 (where enteric γ-Proteobacteria like *E. coli* emerged), *cheD* and the more upstream F7 *cheY* were lost, and the *tar*-like chemoreceptor gene was duplicated. None of the stage 5 genomes retained any genes from the F6 system.

For reference, the species imaged in this study have F7 systems from stage 1 (*V. cholerae* and *M. alcaliphilum)* and stage 2 (*P. aeruginosa* and *S. oneidensis*).

The flagellar motor is controlled by the F6 system in many species with stage 1 or 2 F7 systems (*28–31*), and it is controlled by the F7 system in stage 5 species (the enterics). Our results therefore suggest that control of the motor switched from the ancestral F6 system to the new F7 system in the transition between stages 2 and 3. Less is known about stage 3 and 4 species (β-Proteobacteria), but at least some use flagella and have been reported to be chemotactic(*29*). Based on our results, we would predict that the F7 system controls the flagellar motor in these organisms. The function of the F7 system in stages 1 and 2 remains unclear.

### System inputs: Evolution of the chemoreceptors (Aer2, McpA and Tar)

How did the F7 system take control of the flagellar motor? To address this question we examined both the inputs and outputs of the system. First, we examined the inputs by analyzing the chemoreceptors. In stages 1 and 2, the F7 cluster included both *mc-pA*-like and *aer2*-like receptors, and the F6 system included a *tar-like* receptor. Because nearly all the genomes with stage 1 and 2 F7 systems possessed *mcpA*-like and *aer2*-like chemoreceptors in the gene cluster, both are apparently needed for the (unknown) function of F7 in these organisms. This is puzzling, however, since as described above, we found that McpA-like receptors were non-essential for formation of the F7 array. Suggesting an alternative scenario, a previous study found that F7 McpA-like receptors physically associate with the F6 array(*30*).

To explore the evolutionary history of these two receptors, we identified 130 Aer2-like and 39 McpA-like chemoreceptors from the pool of 166 γ-proteobacterial genomes we used to build the phylogenetic profile in Fig. S2 (some Xanthomonadales contained multiple copies of Aer2-like receptors in their F7 gene clusters, some Shewanellaceae lacked McpA-like receptors and none of the Enteric genomes had either). All 130 identified Aer2-like receptors belonged to the 36H heptad class, consistent with their belonging to F7 systems(*9, 18*). The McpA-like receptors could not be assigned to a heptad class. Inferring the relationships among these receptors with a phylogenetic tree, we found that the evolutionary histories of both Aer2-and McpA-like chemoreceptors are largely congruent with that of the CheABR phylogeny (Fig. S4). More specifically, they recapitulate the split between stage 1 and 2 F7 systems. Proteins often co-evolve when they participate in the same, or codependent, biological functions, so this again suggests that both McpA-like and Aer2-like receptors mediate the ancestral F7 function. This was expected for Aer2-like receptors (which are part of the F7 array), but surprising for McpA-like receptors (which are part of the F6 array). It is unclear what function McpA-like receptors might perform for the ancestral F7 system while physically integrating into the F6 array.

The change in the biological function of the F7 system apparently coincided with the transformation of the *aer2*-like gene into a *tar*-like gene. To investigate this switch we looked further into the domain architecture of the Aer2- and Tar-like MCPs. While their signaling domains are similar (all belong to the 36H heptad class), the rest of their topology differs. As described above, Aer2-like receptors have multiple PAS-HAMP repeats and no periplasmic domain (Fig. 2B). In contrast, 36H F7 Tar-like receptors have a periplasmic N-terminal sensor sandwiched by two transmembrane regions and a single HAMP domain(*31*). Interestingly, this topology is the same as that of the majority of 40H F6 receptors that control flagellar motility in γ-Proteobacteria with stage 1 and 2 F7 systems(*18*). Thus, the F7 system’s takeover of the flagellar motor involved acquisition of a sensory input (an N-terminal periplasmic sensor domain) similar to that of the F6 system that used to control the motor.

### System outputs: Evolution of the response regulator (cheY) and signal termination phosphatase (cheZ)

Finally, we examined the outputs of the system: *cheY* and *cheZ*. We first assigned *cheY* and *cheZ* genes to F7 and F6 systems based on their location in gene clusters (Fig. 3). We observed that organisms with stage 1 or 2 F7 systems possessed *cheY* genes in their F7 and F6 clusters, but only the F6 systems also included *cheZ*. These two genes were retained in stages 3 and 4, even as the F6 cluster was lost. The ancient *cheY* gene from the F7 system was finally lost, in stage 5. To test whether the sequences support this history, we performed a phylogenetic analysis of all *cheY* genes present in the 246 proteobacterial genomes used for the CheABR analysis (Fig. 4). This showed that the “extra” *cheY* genes outside the F7 gene cluster in stage 3 genomes were more closely related to F6 *cheY*s than to the older F7 *cheYs*. This F6-related *cheY* is the same one that appears at the downstream end of the F7 cluster in stage 4, and also the one that remains in stage 5 (as the old F7-related *cheY* gene near the front of the cluster is lost). It was previously shown that the *cheZ* genes that moved into the F7 cluster were descendants of the F6 *cheZ*s (*9*). This supports the notion that as the F7 system took over control of the flagellar motor, it lost its original output and acquired the motor-controlling outputs of the F6 system.

**Figure 4.**
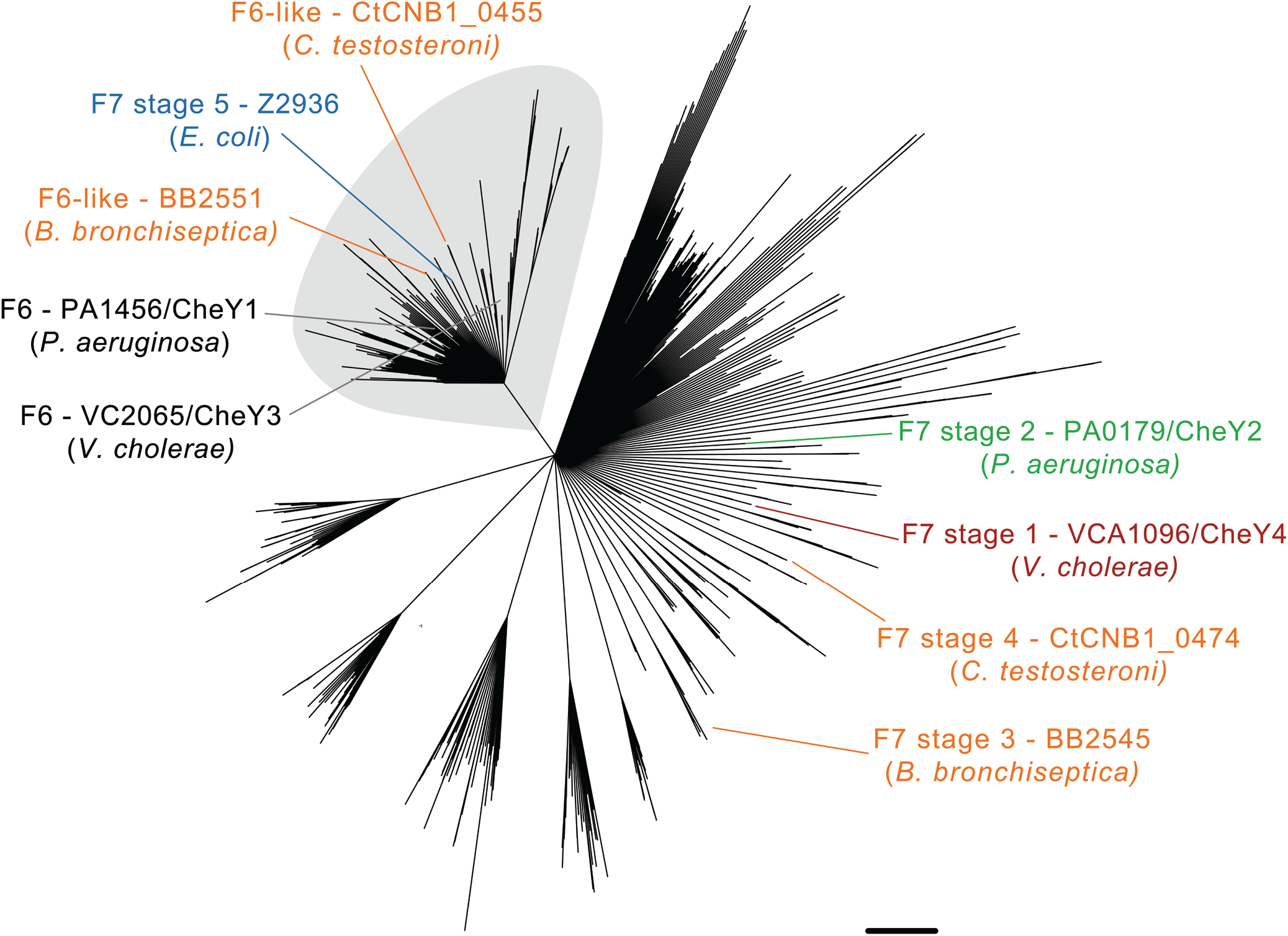
Phylogenetic reconstruction of CheY in selected proteobacterial genomes. Sequences of CheY proteins from F6, F6-like and F7 stage 5 systems cluster in a monophyletic group (grey shade). CheY sequences from the F7 systems of other stages (1, 2, 3 and 4) appear in other clades, indicating that F6-like and F7 stage 5 CheYs are more closely related to CheYs from F6, than F7 systems. Nodes are collapsed on 50% bootstrap support. For each stage, a representative sequence is highlighted, named with the type of its CheY, the locus number / gene name (when annotated) and the name of the organism to which it belongs. These sequence names follow the color code of stages in Fig. 3.

## DISCUSSION

Here, using a combination of cryo-ET and bioinformatics, we have characterized and dissected the evolution of the F7 chemosensory array in γ-Proteobacteria. We find that the ancient F7 system, still present in non-enteric γ-Proteobacteria, took control of the flagellar motor from the F6 system in a series of clear evolutionary steps (Fig. 5A). Thus the well-studied chemosensory model system of *E. coli* is a chimera of two other, more widespread systems: the F6 flagellar-control system and an ancient F7 system of still-unknown biological function. These results provide a striking example of how evolution can repurpose macromolecular complexes for new functions.

**Figure 5.**
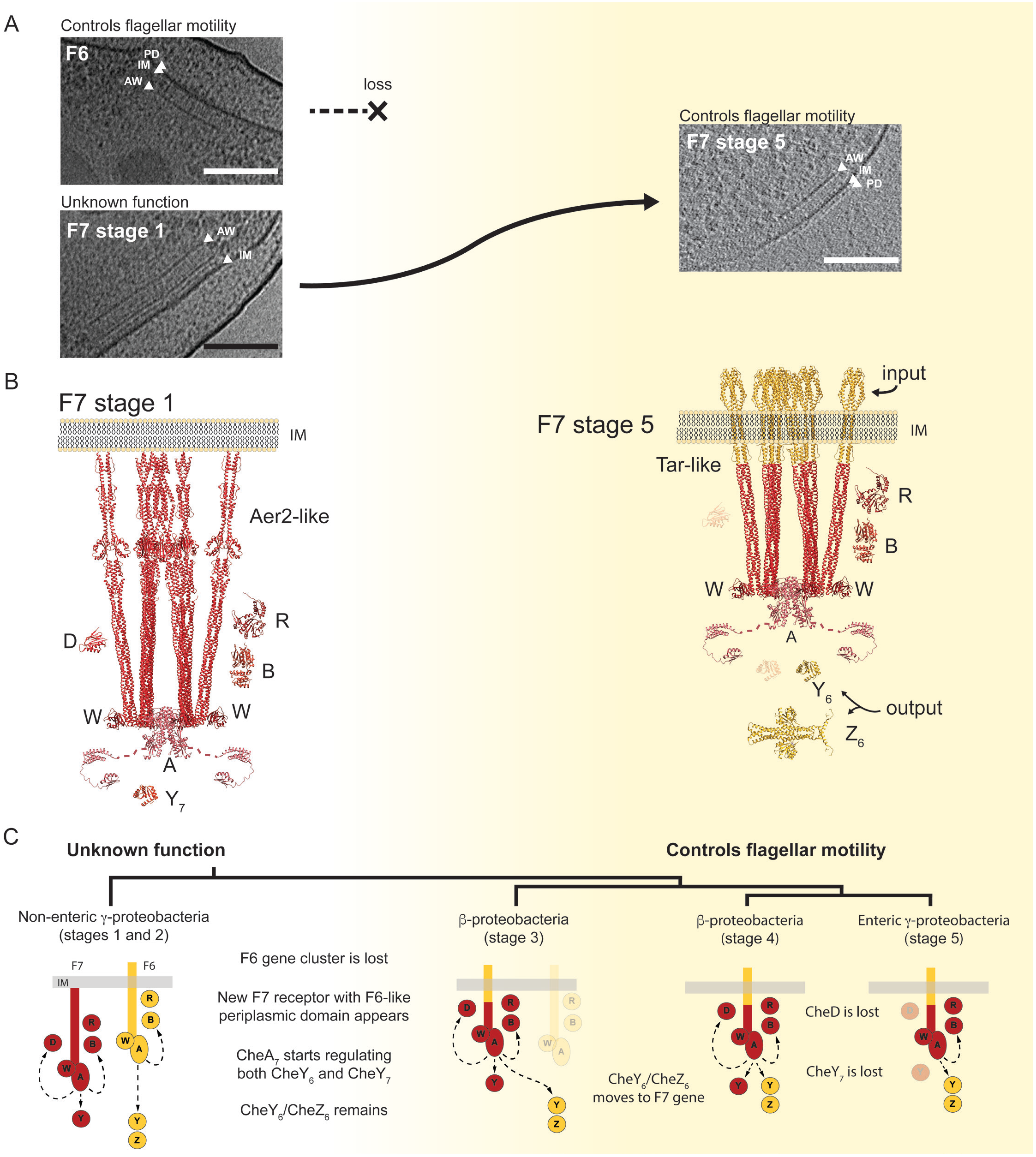
Evolution of the F7 chemosensory array in non-enteric γ-Proteobacteria to acquire F6-like ultrastructure and function. A) Tomographic slices showing F6 and F7 stage 1 chemosensory arrays in the same *P. aeruginosa* cell (left) and an F7 stage 5 chemosensory array in *E. coli* (right). Over the course of evolution, the F6 system is lost and the F7 system evolves similar ultrastructure and function to the F6 system. B) Molecular models of F7 chemosensory arrays in *P. aeruginosa* (left) and *E. coli* (right). Models are colored according to their hypothetical original class: F7 (red) and F6 (yellow). C) Working model of the evolution of the F7 chemosensory system in γ-Proteobacteria and β-Proteobacteria. Scale bars are 50 nm.

We identified four sequential evolutionary steps, each of which produced a stable, modern subtype of chemosensory array. In the step in which flagellar control moved from the F6 to the F7 system, two major evolutionary events were required: (i) the Aer2-like F7 receptor became Tar-like, swapping its input (Fig. 5B); and (ii) the F7 CheA began signaling through the remaining F6 CheY, adding an output. We speculate that this receptor transformation may have occurred via a domain swap that replaced the multiple PAS-HAMP domains of an Aer2-like receptor with the sensor domain of an F6 Tar-like receptor. These changes were accompanied by gradual loss of the remaining F6 components, as well as F7 components no longer needed for its new function (Fig. 5C). Thus, we hypothesize that intermediate stages of the F7 system, present in extant β-Proteobacteria, retain both the older and younger functions.

Our results also shed light on the biological function of CheD. CheD is thought to interact with chemoreceptors in an adaptation mechanism together with CheC and CheY(*32*), but more recent results showed that CheD from *Bacillus subtilis* is able to deamidate chemoreceptors *in vitro* without CheC(*33*). Our results provide two further pieces of evidence supporting the idea that CheD is able to perform a biological role independently of CheC. First, the ancestral F7 system included *cheD* but not *cheC*. Second, *cheD* co-evolved with the ancestral F7 *cheY* (it was lost in the same evolutionary step), pointing to a functional link.

Imaging four γ-Proteobacteria with both F6 and F7 systems by cryo-ET, we observed that the arrays formed by different chemosensory systems were both separate and structurally distinct. This is consistent with previous studies showing physical separation of the arrays from different chemosensory gene clusters in *V. cholerae* (F6 and F9 systems (*18*)) and *P. aeruginosa* (F6 and F7 systems(*30*)). It is also consistent with experiments in *E. coli* showing that engineered chemoreceptors with additional heptads cannot form arrays with shorter, native chemoreceptors, likely because of a large physical mismatch in the CheA/CheW layers(*34*).

In all cases, we observed that the F7 arrays in non-enteric γ-Proteobacteria were membrane-associated, but it remains unclear how this is achieved. In other arrays, N-terminal hydrophobic patches of chemoreceptors mediate membrane insertion. Aer2-like receptors in *V. cholerae, P. aeruginosa* and *S. oneidensis*, however, lack any predicted transmembrane regions. The *M. alcaliphilum* Aer2-like receptor has two small fragments of transmembrane regions (10 and 14 residues), but these are likely too short to attach the receptor to the membrane. One possibility is that the receptors are post-translationally modified for membrane attachment. Another possibility is that another protein serves as a membrane anchor. Our work ruled out one such candidate – the McpA receptor in the same gene cluster; *ΔmcpA* F7 arrays were still attached to the membrane.

One of the most striking features of the F7 arrays in non-enteric γ-Proteobacteria was the presence of additional density layers between the CheA/CheW baseplate and the IM. Based on our homology models of the receptors, we propose that these layers correspond to domain features (Fig. 2B). The L2 layer matched the PAS domain present in Aer2-like receptors in all four species. The fainter (possibly less-ordered) L3 layer in *V. cholerae* and *S. oneidensis* matched the additional PAS domain in the Aer2-like receptors from these species. This suggests that PAS domains might mediate in-tra- and inter-trimer interactions, potentially contributing to cooperativity in the signaling array. The L1 layer matched the junction between the HAMP and signaling domains, which is puzzling since that region of the chemoreceptor structure is predicted to have low molecular density(*35*). It is unlikely that this density is produced by another protein, for example CheR, which is known to bind the chemoreceptor in that area. However, the abundance of this protein appears to be too low to generate a visible density layer, at least in *E. coli* and *B.subtilis* where the chemosensory protein stoichiometry has been determined (*36, 37*). Furthermore, previous cryo-ET of *in vitro* preparations containing only *E. coli* CheA, CheW and Tsr showed a similar layer in that region, suggesting one or more of these proteins alone is responsible for the L1 layer(*38*).

Another mystery is the function of the F7 chemosensory array in non-enteric γ-Proteobacteria. Flagellar motility in these organisms is controlled by the F6 chemosensory system(*39–41*), which is expressed under a variety of conditions. In contrast, the *P. aeruginosa* and *V. cholerae* F7 system is only expressed when cells are grown in stressful conditions such as into late stationary phase, induced by the stress-related sigma factor RpoS(*30, 42, 43*). Expression of the F7 system in different conditions has not been studied in *S. oneidensis* or *M. alcaliphilum*, but both organisms live in unique and challenging environments which may be poorly mimicked by laboratory growth; *S. oneidensis* is a facultative anaerobe adapted to changing environments (*44*) and *M. alcaliphilum* is a haloalkaliphilic methanotroph(*21*). While we did not test different growth conditions for *M. alcaliphilum*, we did observe that formation of F7 arrays in *S. oneidensis* was dependent on culture conditions. Another clue is that both *P. aeruginosa* and *V. cholerae* are capable of sensing oxygen, which binds to the PAS-heme domains of Aer2 receptors to activate signaling(*24, 45*). We therefore favor the working model that the older F7 systems are part of an emergency response system activated by stress conditions, perhaps related to the availability of oxygen. The McpA receptor may also be an important mediator of this response. McpA has no sensory domain, but has been implicated in taxis toward trichloroethylene(*46*). A previous study in *P. aeruginosa* showed that McpA physically co-localizes with F6 system proteins(*30*). Here we find that despite being part of the F6 system, McpA co-evolved with the F7 system, suggesting that McpA may bridge the two systems to provide additional inputs to the flagellar control system in response to stress.

## ACKNOWLEDGMENTS

The authors wish to thank Drs. Zhiheng Yu, Jason de la Cruz, Chuan Hong and Rick Huang for microscopy support at HHMI Janelia Farms; Dr. Mohamed Y. El-Naggar for insights into the culturing of *S. oneidensis*; Dr. Kristin Wuichet for discussions about the evolutionary interpretation of the bioinformatics data and Dr. Keith Cassidy for discussions on homology model building. We also thank Dr. Catherine M. Oikonomou for helpful discussion and suggestions on the manuscript. Cryo-EM work was done in the Beckman Resource Center for Transmission Electron Microscopy at Caltech.

## Funding

This work was supported by the John Templeton Foundation as part of the Boundaries of Life Initiative (grants 51250 & 60973), NIGMS grant R35 GM122588 (to GJJ), NIGMS grant R01 GM108655 (to KJW), the Max-Planck-Gesellschaft (to SR), and the Air Force Office of Scientific Research Presidential Early Career Award for Scientists and Engineers grant FA955014-1-0294 (SP). *P. aeruginosa* strains were acquired from the transposon mutant collection that was made possible by NIH grant P30 DK089507.

## Author contributions

Conceptualization: DRO, AB and GJJ; Formal analysis: DRO; Funding acquisition: KJW, MGK, SR, AB and GJJ; Investigation: DRO, PS, PM, AK, SC, KJW, SP, SAC, RK and AB; Methodology: DRO, PS, PM, KJW, SP, DAC, MGK, SR and AB. Project administration: GJJ; Software: DRO; Supervision: MGK, SR, AB and GJJ. Validation: DRO, PS, PM, AK, SC, KJW, SP, SAC, RK and AB; Visualization: DRO, PS, SP and AB; Writing – original draft: DRO, PS, AB, GJJ. Writing – review and editing: DRO, PS, KJW, SP, DAC, MGK, SR, AB and GJJ.

## Competing interests

Authors declare no competing interests;

## Data and mate-rials availability

1D electron density profile tool is available on node package manager (npm): https://www.npmjs.com/package/sideview-profile-average. The script to visualize the 1D electron density profile and calculate uncertainties is available on Observable HQ: https://observablehq.com/d/6399ff6b27e2465b. Regular Architecture code is available on npm: https://www.npmjs.com/package/regarch. Tomograms are available in the Electron Tomography Database – Caltech at https://etdb.caltech.edu. Phylogenetic trees and homology models are available as Supplementary Datasets.

## SUPPLEMENTARY MATERIALS

Materials and Methods

Tables S1-S9

Figs S1-S4

References (47-76)

Datasets S1-S2

## Notes

#### Summary of Updates

fix typo in author's list

