## Supplementary Materials for "Repurposing a macromolecular machine: Architecture and evolution of the F7 chemosensory system"

**This PDF file includes:**

Materials and Methods
Figs. S1 to S4
Tables S1 to S9
Captions for Datasets S1 to S2

**Other Supplementary Materials for this manuscript include the following:**

Datasets S1 to S2

### Materials and Methods

#### Strains and growth conditions

*Vibrio cholerae*

Strain list and construction:
Wild-type: *Vibrio cholerae* C6706
Strain PM6: *Δvca1088*
Strain PM7: *Δvca1093*, *Δvca1094*, *Δvca1095* (ΔF7)
Strain PM18: *Δvca1092*

*V. cholerae* deletion strains were generated using standard allele exchange (*47*) with the plasmids listed below.

Plasmid for deletion of *vca1093*, *vca1094* and *vca1095*:
Plasmid pPM045 was constructed by PCR amplification of the up- and down-stream regions of *vca1093* and *vca1095*, respectively. PCR1 was performed with primers ccccctctagaaattggctaatccctcctaaactc / aatcttgcgcagttgttccatatc and C6706 chromosomal DNA as template. PCR2 was performed with primers gatatggaacaactgcgcaagatt cgcttaagcaccactgccgaa / ccccctctagacatcatcaaattcgtcgtcatgc and C6706 chromosomal DNA as template. A third PCR was then performed using primers ccccctctagaaattggctaatccctcctaaactc / ccccctctagacatcatcaaattcgtcgtcatgc and PCR1 and PCR2 as template. The product from PCR3 was then digested with XbaI and ligated into the equivalent site of plasmid pCVD442 (*47*) resulting in plasmid pPM045.

Plasmid for deletion of *vca1092*
Plasmid pPM051 was constructed by PCR amplification of the up- and down-stream regions of *vca1092*. PCR1 was performed with primers ccccctctagaacggttgtcttgatcttgagtgc / aacaaactggggcacaacctg and C6706 chromosomal DNA as template. PCR2 was performed with primers caggttgtgccccagtttgttatgcataaagcaccgataaatcagg/ ccccctctagaattgccttgctgatctttgaccc and C6706 chromosomal DNA as template. A third PCR was then performed using primers Ccccc tctaga acggttgtcttgatcttgagtgc / ccccctctagaattgccttgctgatctttgaccc and PCR1 and PCR2 as template. The product from PCR3 was then digested with XbaI and ligated into the equivalent site of plasmid pCVD442 resulting in plasmid pPM051.

Plasmid for deletion of *vca1088*
Plasmid pSR1228 was constructed by PCR amplification of the up- and down-stream regions of *vca1088*. PCR1 was performed with primers ccccctctagaaagccaatgtagggtttgtgcag / tatcgccgttattttgtgttttctcg and C6706 chromosomal DNA as template. PCR2 was performed with primers cgagaaaacacaaaataacggcgataaaccgtgggggattggctg / ccccctctagatgcgataatgtgcctgtactttg and C6706 chromosomal DNA as template. A third PCR was then performed using primers ccccctctagaaagccaatgtagggtttgtgcag / ccccctctagatgcgataatgtgcctgtactttg and PCR1 and PCR2 as template. The product from PCR3 was then digested with XbaI and ligated into the equivalent site of plasmid pCVD442 resulting in plasmid pSR1228.

During plasmid and strain construction, *V. cholerae* and *E. coli* were grown at 37°C in LB medium or on LB agar plates containing antibiotics in the following concentrations: 200 mg/mL streptomycin; 50 mg/mL kanamycin; 100 mg/mL ampicillin; 50 mg/mL carbenicillin; and 20 mg/mL chloramphenicol for *E. coli*, and 5 mg/mL for *V. cholerae*.

For cryo-ET microscopy *V. cholerae* cells were grown as described previously (*16*): after growth in LB medium for 24 hours at 30°C with shaking, 150 µl cell suspension was diluted into 2 ml Ca-HEPES buffer and grown for an additional 16 hours with shaking at 30°C.

*Pseudomonas aeruginosa*

The *P. aeruginosa* mutant strains imaged in this study were acquired from the transposon mutant collection from the University of Washington, Table S4. Wild-type *P. aeruginosa* PAO1 and a PAO1 *aer2* deletion mutant [PAO1047](*30*) were imaged in this study. Cells were grown in MOPS-based nitrogen starvation medium for ~24 hours at 30°C with shaking. MOPS-based minimal medium limited nitrogen(*48*): 43mM NaCl, 50mM MOPS (from 1M stock of MOPS/NaMOPS pH 7.2), 40 mM Sodium Succinate, 1mM $MgSO_4$, 2.2mM KCl, 0.1mM CaCl, 10µM $FeNH_4SO_4*7H_20$, 1 mM $NH_4Cl$, 1.25mM $NaH_2PO_4$.

*Shewanella oneidensis*

For imaging the chemosensory systems, *Shewanella oneidensis* MR-1 wild-type cells were cultured using continuous flow bioreactors (chemostats) or batch cultures as previously described(*20*).

*Methylomicrobium alcaliphilum*

For imaging intracellular structures wild type *M. alcaliphilum* $20Z^R$ cells were grown in a modified nitrate mineral salts medium(*49*) with a final pH of 9.0 consisting of: 9.9mM $KNO_3$, 0.8mM $MgSO_4$ x 7 $H_2O$, 13.6µM $CaCl_2$ x 2 $H_2O$, 0.5M $g*L^{-1}$ NaCl, 2mM $KH_2PO_4$, 2mM $Na_2HPO_4$, 22.5mM $NaHCO_3$, 2.5mM $Na_2CO_3$ along with trace elements 13.4 µM $Na_2EDTA$, 7.2µM $FeSO_4$ x 7 $H_2O$, 4.8µM $CuSO_4$ x 5 $H_2O$, 1 µM $ZnSO_4$ x7 $H_2O$, 0.9 µM $Na_2O_4W$ x $2H_2O$, 0.8 µM $CoCl_2$ x 6 $H_2O$, 0.5µM $H_3BO_3$, 0.2 µM $MnCl_2$ x 4 $H_2O$, 0.2µM $NiCl_2$ x 6 $H_2O$, 0.2µM $Na_2MoO_4$ x 2 $H_2O$. Cultures were grown at 30°C in septated bottles containing 20% $CH_4$ headspace. Cell samples for cryo-ET preparations were taken at mid-exponential phase, at a cell density of $OD_{600}$ = 0.5.

**Electron cryotomography**

Cells were prepared for electron cryotomography as described previously(*50*) . Images were collected using either an FEI Polara 300 keV field emission gun microscope or an FEI TITAN Krios 300 keV field emission gun microscope with lens aberration correction (FEI Hillsboro, OR). Both microscopes were equipped with Gatan imaging filers and 'K2 summit' counting electron detector cameras (Gatan, Pleasanton, CA). The data collection software used to collect the tilt series was UCSFtomo (*51*). The cumulative electron dose was 160 $e^-/A^2$ or less for each individual tilt series. CTF correction, frame alignment and SIRT reconstruction were done using the IMOD software package (*52*). The tomograms used in this study are available in the Electron Tomography Database – Caltech (*53*) and their identifiers can be found in Table S5.

**1D electron density profiles**

To measure the distance between the inner membrane (IM) and the CheA/CheW base plate, we used a custom script written in Node.js that takes as input the tomogram and the model points that delineate the inner membrane using 3dmod. The script calculates the average pixel value in profiles running perpendicular to the model points but in the same plane as the model points were collected. These averaged profiles are output in a JSON formatted file. The script and instructions for installation and use are available in a GitLab repository at https://gitlab.com/daviortega/sideview-profile-average. To visualize the profile we used the ObservableHQ notebook located at https://beta.observablehq.com/@daviortega/generic-notebook-to-analyse-1d-averaged-electron-density-p. For each profile, we measure the distance between the peaks corresponding to the electron density of the IM, the CheA/CheW baseplate and the intermediate layers when possible, in pixels. The measurement uncertainty was estimated as the expanded uncertainty using k = 2 as a coverage factor propagated from the uncertainty in determining the center of each peak in pixels, as recommended by (*54*). The values reported in nanometers were calculated by multiplying the measurement and uncertainty by the pixel size for each tomogram.

**Bioinformatics resources and software packages**
All sequences in this study were collected in the MiST database (*55*), domain architecture predictions from PFAM(*56*) were selected from SeqDepot (*57*), and 3D atomic models were taken from the Protein Data Bank (PDB) (*58*). Domain architecture prediction was performed with CD-VIST (http://cdvist.zhulinlab.org/)(*25*). We used CD-HIT v4.6 to reduce redundancy in unaligned sequences(*59*). Multiple sequence alignments were performed with the algorithm L-INS-I from the package MAFFT v7.305b (*60*). We used Gblocks v 0.91b (*61*) to eliminate poorly aligned columns in multiple sequence alignments. To perform sequence alignments with structural information we used STRAT from the MultiSeq (*62*) tool for VMD v1.9.3 (*63*) which in turn was used to visualize and manipulate 3D structures. Homology modeling was performed using MODELLER v9.17 (*64*). Secondary structure predictions were performed with JNet Structure Predictions(*65*) in the Jalview(*66*) software, which was also used to visualize multiple sequence alignments. Similarity searches for sequences were conducted using BLAST v2.7.1+ (*67*) and HMMER v3.1b2 (*68*). Phylogenetic reconstructions were performed using RAxML v8.2.10(*69*). To collapse branches with low support in phylogenetic trees we used TreeCollapseCL4 (*70*). Tomograms and model points were manipulated using 3dmod v4.9.9 (*52*).

**Protein domain architecture prediction**
The domain architecture of the C-termini of chemoreceptors are poorly conserved among members of this protein family(*71*). Further, protein domains commonly appearing in this region, such as HAMP(*72*) and PAS(*26*), are so diverse that in several instances predictive models have difficulty identifying them. To address this problem we used CD-VIST on the two Aer2-like receptors without known domain architectures from *S. oneidensis* and *M. alcaliphilum* with TMHMM prediction, skipping HMMER3 and RPSBLAST steps, but adding three consecutive HHSEARCH steps against the PDB database with HHBLITS using uniclust30 at different thresholds for minimum probability: 60%, 40% and 20%. Analyzing the CD-VIST domain coverage we predicted that *S. oneidensis*'s Aer2-like receptor has a PAS-PAS-HAMP-HAMP-MCPsignal domain architecture, similar to *V. cholerae* and that the *M. alcaliphilum*'s Aer2-like receptor has a HAMP-PAS-HAMP-HAMP-MCPsignal domain architecture. We further enhanced our confidence in this predictions by aligning the Aer2-like sequences to the sequence of the templates used to produce the homology models.

**Homology modeling**
To build homology models for the Aer2-like receptors in *V. cholerae*, *P. aeruginosa*, *S. oneidensis* and *M. alcaliphilum* we used several crystal structures available in the Protein Data Bank (PDB), Table S6. The files used in this process are described in Table S7 and can be found in the Supplementary Dataset S1.

First we built a homology model with two HAMPs followed by the MCPSignal domain that we name 2H+S. For that we used the structures 3ZX6 and the second HAMP of 4I3M to form a chimeric template. We manually aligned the structures of the templates against the Aer2 in *P. aeruginosa* (PA1076) and performed a multiple sequence alignment using L-INS_I and MultiSeq. To construct the homology model of this structure, we use MODELLER with the following parameters: a.library_schedule = autosched.slow, a.max_var_iterations = 1000, a.repeat_optimization = 100 and a.max_molpdf = 1e6. To make sure that the connection between both HAMPs remained the same, we added a restraint in both chains A and B from residues 359 to 385. We built 100 homology models with these parameters and chose the one with the lowest DOPE score.

Next, to add a PAS domain to this structure, we used the 3VOL and 4HI4 structures. First we aligned chain B of 3VOL with the 2H+S homology model produced in the previous step. We noticed that this alignment produced clashes between the PAS domains. To overcome this obstacle, we used chains B and D in the 4HI4 structure as a model for the dimerization of the two PAS domains. We aligned chain B of 4HI4 to the 3VOL structure using the residues QWTDRT and then manually manipulated the dimer of PAS to be positioned in line with the 2H+S model to build the next homology model: P+2H+S. Sequence alignment was performed as described before against the sequence of Aer2 in *P. aeruginosa* (PA1076). This homology model was used as the basis of the complete homology models of all the Aer2-like receptors.

To build the homology model of Aer2 in *P. aeruginosa* (PA1076), we used the P+2H+S model together with the 4I3M structure. For that we manually aligned the structures to build the template. However, there is a 13 residue region unresolved in both structures (R156 – G169) but predicted to be alpha helical. We assume that these two structures then are around 2.2 nm apart and took that into consideration while positioning the structures. Finally the homology model was built using MODELLER with the parameters described above and with a restraint to force alpha helical conformation between residues 140 to 181.

To build the homology model of the Aer2-like receptor in *V. cholerae* (VCA1092) we used the P+2H+S model together with 4HI4. The sequences of the templates and VCA1092 aligned pretty well with only a minor gap in the residues ELLRD, also predicted to be alpha helical. We aligned the end of chain B of the already aligned 4HI4 used in the P+2H+S model to the beginning of chain A of P+2H+P using STAMP and manually adjusted the position of the structures using VMD. The homology model was constructed with MODELLER and we imposed a restraint to force alpha helical conformation between residues 21 to 43 (C terminal) and 151 to 171 (unresolved gap).

To build the homology model of the Aer2-like receptor in *S. oneidensis* (SO_2123) we used the VCA1092 model since they have the same domain architecture. The sequences of the templates and SO_2123 also aligned pretty well with only a minor gap in the residues ESIDA, also predicted to be alpha helical. The C-terminus of the sequence is also predicted to be alpha helical up to the residue PHE7. We aligned the end of chain B of the already aligned 4HI4 used in the P+2H+S model to the beginning of chain A of P+2H+P using STAMP and manually adjusted the position the structures using VMD. The homology modeling was performed with MODELLER and we imposed a restraint to force alpha helical conformation between residues 21 to 43 (C terminal) and 151 to 171 (unresolved gap).

To build the homology model of the Aer2-like receptor in *M. alcaliphilum* (MEALZ_2872) we used the P+2H+S model and the 4I3M structure. To find out which of the 3 HAMPs in the 4I3M structure is most closely related to the C-terminal HAMP of MEALZ_2872 we used BLAST to find HAMP sequences in the *Pseudomonas* group similar to each of the HAMPs in the 4I3M and to the C-terminal HAMP of MEALZ_2872. We aligned the sequences using L-INS-I and perform a phylogenetic reconstruction using RAxML with -m PROTGAMMAIAUTO -p 1234555 -x 9876545 -f a -N 100 as parameters. Tree nodes were collapsed to a certainty score of 50. The phylogenetic analysis showed that the C-terminal HAMP of MEALZ_2872 is closely related to the second HAMP of 4IM3. We truncated the 4IM3 structure to contain only the second HAMP and aligned an extended helix connecting to the third HAMP with the PAS domain of the P+2H+S model. We used this alignment to place the HAMP at the right position and deleted the extended helix. These structures were used as a template for the MEALZ_2872 homology model built with MODELLER as described above and with restraints to force alpha helical conformation in residues 194 to 216, 255 to 270 and 721 to 728.

**Chemotaxis system classification**

Relevant protein sequences of chemotaxis components were classified using HMMER and the hidden Markov models previously published (*9*). The model with highest score was used to assign chemotaxis components to classes.

**F7 system identification in γ-Proteobacteria.**

To estimate how widespread F7 systems are in γ-Proteobacteria, we randomly picked 310 genomes from γ-Protebacteria from MiST. From those, we selected the CheA protein sequences and then classified them using HMMs provided by the authors of (*9*). The CheA proteins classified as F7 systems belonged to 176 genomes. Table S8 list all 310 genomes and marks the presence of the F7 systems in the 176 genomes.

**Phylogenetic tree of F7 systems in Proteobacteria**

To build a tree of the F7 and F8 systems in Proteobacteria we used a concatenated alignment of the protein sequences of CheA, CheB and CheR, as previously described (*9*). We first collected every CheA belonging to these two classes from 1152 Proteobacteria genomes in MiST (547 from F7 class and 168 from F8 class) and used CD-HIT to eliminate redundancy at the 85% identity level (201 from F7 class and 119 from F8 class). To find CheB and CheR proteins that confidently function with the selected CheAs, we searched for genes that code for these proteins in the range of 10 genes upstream and downstream from each *cheA* gene. Conflicts of multiple or missing *cheB*, *cheR*, or *cheA* genes within that range were manually resolved or the system was removed from the dataset. At this stage the dataset contained 272 protein sequences of CheAs, CheBs and CheRs. We aligned each protein individually with L-INS-I from MAFFT. We used Jalview to examine the alignment and removed 10 sequences for not being complete genes and re-aligned the sequences with L-INS-I. The final dataset had 262 sequences from 246 genomes. For each protein family, we used Gblocks to remove alignment positions with low information. The Gblock parameters were: b3=8 -b4=10 b5=h. The resulting alignments of the protein sequences of CheA, CheB and CheR were concatenated into a single alignment with 698 columns. We used RAxML with parameters -m PROTGAMMAIAUTO -f d -d -N 25 with different seeds 10 times and 3 partitions set to evolutionary model AUTO with boundaries 1-312, 313-559 and 560-698 to accommodate possible differences in the evolutionary models selected for CheA, CheB and CheR sequences. We selected the tree with best maximum likelihood score. We also ran 1000 rapid bootstrap on the same alignment with the parameters -m PROTGAMMAIAUTO -p 1234555 -x 9876545 -f a -N 1000. We mapped these bootstrap values to the best tree and used TreeCollapseCL4 to collapse nodes with less than 50% uncertainty to polytomies. We also mapped the CheA gene neighborhoods (15 genes up and downstream) to the CheABR tree using custom scripts written in Python to produce Fig. S1. BLAST all vs. all to all CheAs and selected neighboring genes was used to loosely define homologous sets of proteins with at least 10E-40 E-value and query coverage of 95% to any member of the set. As an exception to this rule, the anti-signa factor antagonists were selected with the threshold of 1E-5 and query coverage of 50%. Homologs of relevant proteins are highlighted in different colors. We manually selected representatives of relevant genes neighboring CheA for major branches relevant to this study for display in Fig. 3.

**Phylogenetic profiles of F6 and F7 systems**

We first selected the genomes of the organisms we imaged: *Methylomicrobium alcaliphilum*, *20Z*, *Pseudomonas aeruginosa PAO1*, *Shewanella oneidensis MR-1*, *Vibrio cholerae O1 biovar El Tor str. N1696*. In order to perform phylogenetic profiling of the chemotaxis systems in γ-Proteobacteria, we added 162 genomes from this class and 10 genomes from β-Proteobacteria as an outgroup, for a total of 176 genomes (Table S9). The number of selected genomes is coincidently the same as the number of genomes from γ-Proteobacteria with F7 systems described above but only a fraction of genomes are present in both sets. To build the organism tree we used the same procedure as described in (*73*) with the difference that the final concatenated alignment served as an input to RAxML to generate 164 inferences with the parameters -m PROTGAMMAIAUTO -p 12345 -f d -d -N 164. Chemotaxis proteins from these genomes were classified as described above and mapped onto the organism tree to produce Fig. S2.

**Domain architecture prediction of chemoreceptors present in stage 3 and 4 F7 systems**

We selected the protein sequences of chemoreceptors present in the gene neighborhood used to build Fig. S1 and use CDVIST to predict the domain architecture using TMHMM, HMMER3 against Pfam 30.0 database. The results are shown in Fig. S3.

**Identification of Aer2-like and McpA-like receptors**

We first collected all 3389 chemoreceptors from the 176 genomes used to build the phylogenetic profiles. We defined a protein as a chemoreceptor if it contained the MCPsignal PFAM domain. Then we grouped them in clusters of orthologous groups using the same technique described in

(*18*). To pick Aer2-like receptors we used an E-value of 1E-135 and selected all 144 receptors present in the same group as the Aer2 (PA1076) from *P. aeruginosa*. From those, we removed 6 receptors from the β-Proteobacteria outgroup, 5 that were not classified as 36H receptors and 3 other sequences that did not seem to align well with the group. The final set of Aer2-like receptors had 130 Aer2-like receptors and was aligned using L-INS-I and manually inspected with Jalview. Chemoreceptor families and subfamilies are prone to have diverse C-terminal domain architectures so following the procedure in (*71*) we manually trimmed the sequences to only contain the regions common to all receptors. This final alignment was used to build a phylogenetic tree with RAxML. We built 200 independent inferences with parameters -m PROTGAMMAILG -p 1234555 -f d -d -N 200 and 1000 rapid bootstrap trees with -m PROTGAMMAILG -p 1234555 -x 9876545 -f a -N 1000. Bootstrap scores were mapped to the tree with best maximum likelihood from the 200 independent inferences. Nodes were collapsed to polytomies at 50% uncertainty using TreeCollapseCL4. The same procedure was executed to make the tree of McpA-like receptors but with an E-value threshold of 1E-30. The McpA-like cluster was defined as the one containing McpA from *P. aeruginosa* (PA0180). There were 40 McpAs in the final dataset. Both trees are displayed in Fig. S4.

**Phylogenetic tree of CheY**
The CheY protein comprises a single domain, known in the PFAM database as Response Regulator (Response_reg). However, this domain appears in several other proteins as well. In order to select proteins with one and only one response regulator domain, we collected the domain architecture information from PFAM v30 and predicted transmembrane regions by TMHMM from SeqDepot for all sequences from the 246 genomes and used Regular Architecture (https://www.npmjs.com/package/regarch) to filter only single CheY domains with the following rule:

```
{
  "patterns": [
    {
      "npos": [
        {
          "count": "{0}"
          "name": "TM",
          "resource": "tmhmm2",
        }
      ],
      "pos": [
        {
          "name": "^",
          "resource": "regarch"
        },
        {
          "name": "Response_reg",
          "resource": "pfam30"
        },
        {
```

```
                "name": "$",
                "resource": "regarch"
            }
        ]
    }
  ]
}
```
This pattern selected 4941 sequences. We then used the same clustering techniques described for Aer2-like and McpA-like receptors with an E-value threshold of 10E-30 and selected the largest group, with 1394 sequences. This group contains the known CheYs of the model organisms in this study and others. We aligned this dataset with L-INS-I and manually removed 3 sequences that were highly divergent using Jalview. We built the tree with 500 rapid bootstraps with RAxML and searched for the best tree of this set with the parameters -m PROTGAMMALG -p 1234555 -x 9876545 -f a -N 500. Finally we collapsed nodes with less than 50% support into polytomies using TreeCollapseCL4 to produce Fig. 4.

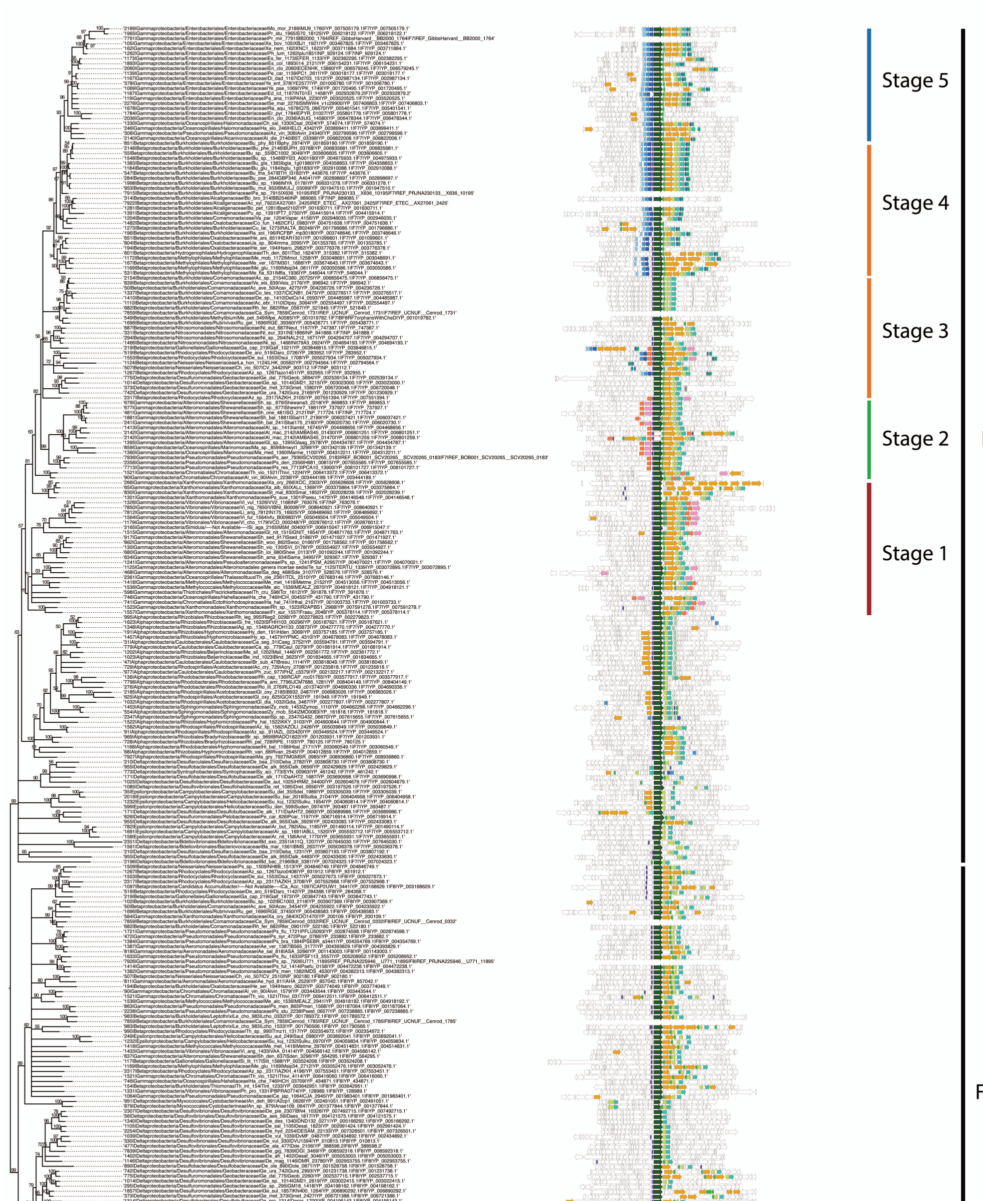


**Fig. S1.**

Phylogeny of CheA, CheB and CheR concatenated alignments of F7 and F8 systems and gene neighborhood of 15 genes up and downstream from CheA.

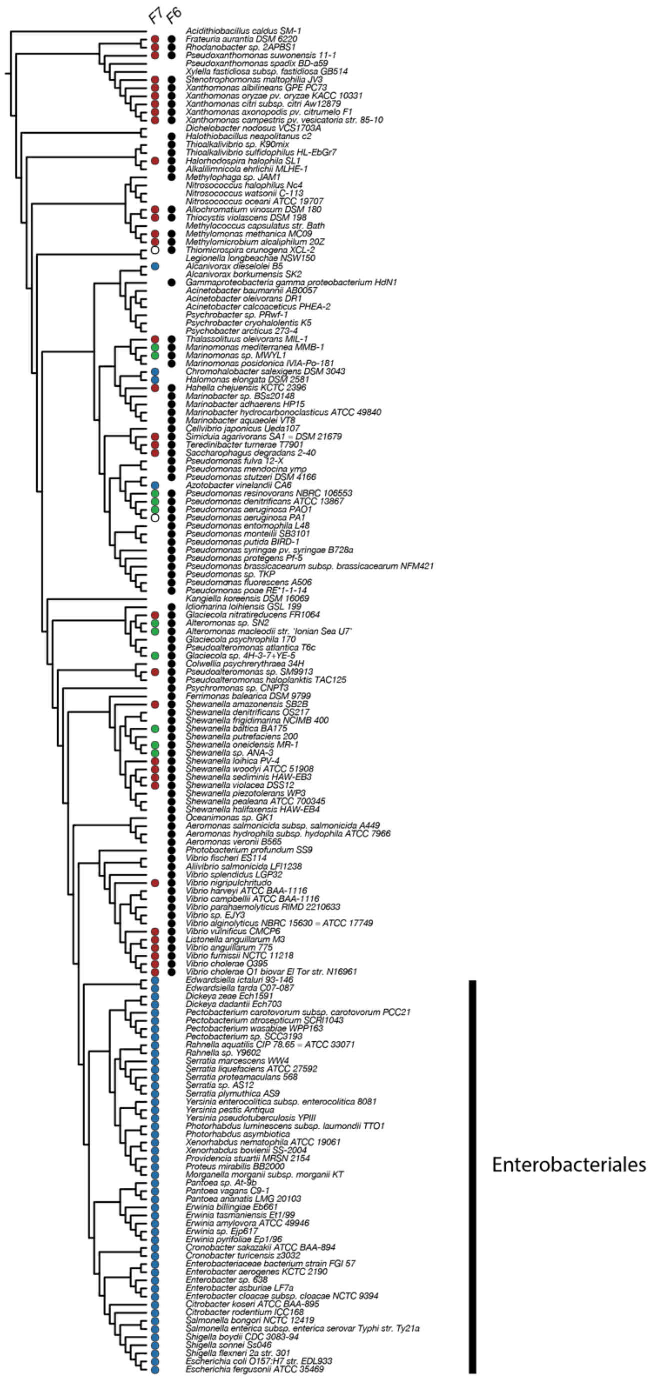


**Fig. S2.**

Phylogenetic profile of the F7 and F6 systems in γ-Proteobacteria shows that only organisms with stage 1 (red) and from stage 2 (green) has F6 systems but not from stage 5 (blue). Note that the distribution of stage 1and stage 2 are mixed in the non-enteric group. Genomes with empty circles were genomes included in this part of the research but not in the analysis of classifying the F7 systems.

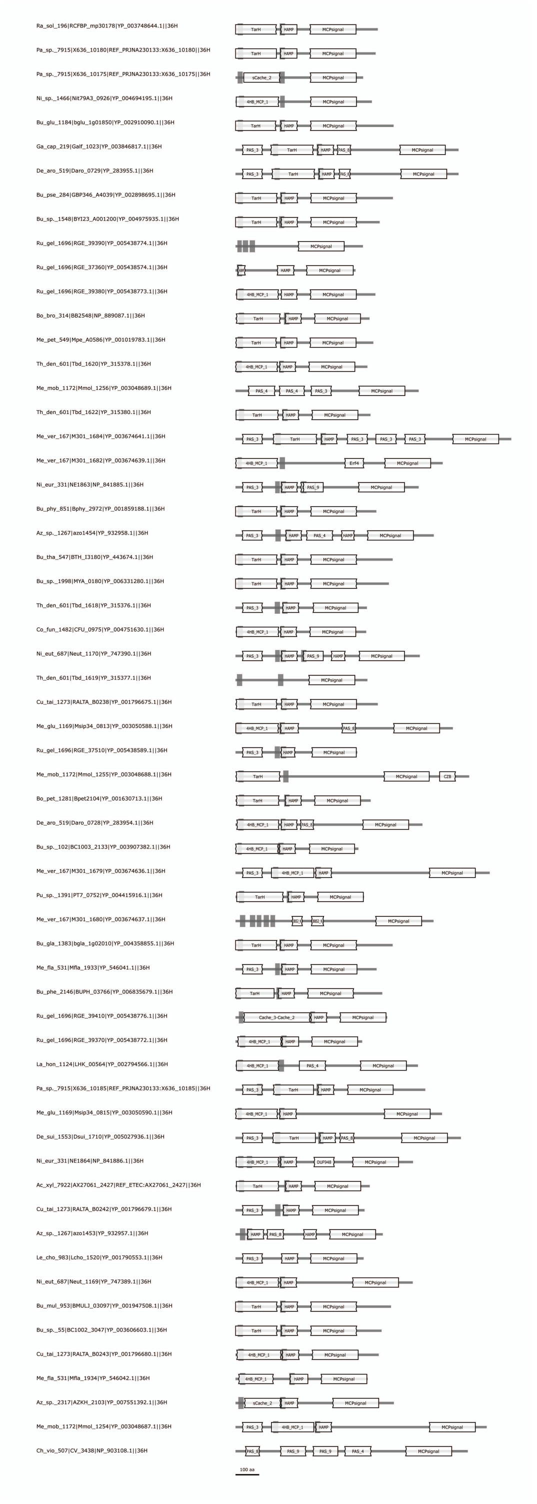

**Fig. S3.**

Protein domain architecture of chemoreceptors in the gene cluster of stage 3 and 4 of F7 systems shows the high incidence of receptors with transmembrane regions and periplasmic sensory domains (TarH, Cache superfamily, 4HB_MCP_1).

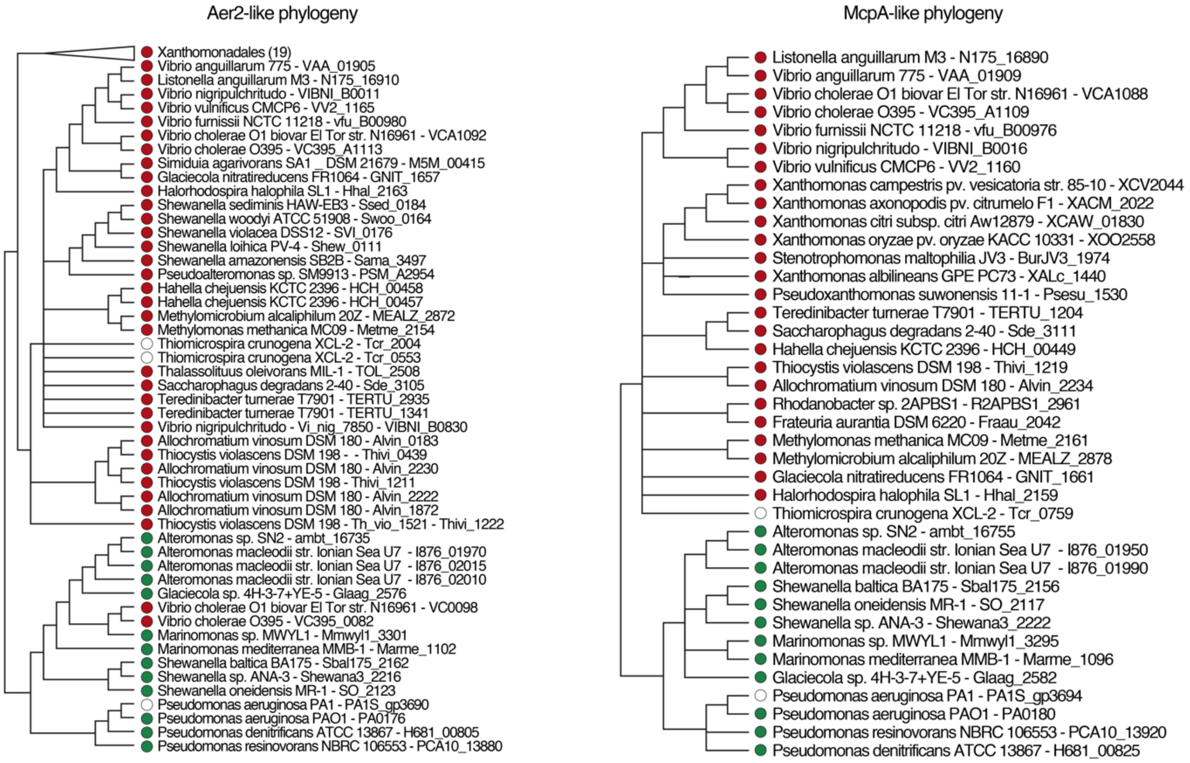


**Fig. S4.**

Phylogenetic tree of Aer2-like receptors and McpA-like receptors. The tags in the tips are built using the name of the organism and the locus of the receptor. Tips with red dots belong to chemoreceptors from stage 1 and green to stage 2. The only exception to the monophyletic distribution of Aer2-like receptors were in *V. cholera*, where an addition to the Aer2 homolog, a second, orphan 36H receptor (VC0098) was likely introduced by a recent lateral gene transfer from *Marinomonas*. Genomes with empty circles were genomes included in this part of the research but not in the analysis of classifying the F7 systems.

**Table S1.**

Presence and absence of the chemosensory arrays in imaged strains of *V. cholerae*, *P. aeruginosa*, *S. oneidensis*, and *M. alcaliphilum*.

| | Imaged cell poles | Short array | Tall array |
|---|---|---|---|
| ***Vibrio cholerae*** | | | |
| Wild-type C6706 | 29 | 20 | 7 |
| *Δmcp (VCA1088)* | 20 | 19 | 5 |
| *ΔF7 cheW, cheW, cheA (VCA1093, VCA1094, VCA1095)* | 29 | 18 | 0 |
| *Δmcp (Aer2/VCA1092)* | 27 | 24 | 0 |
| ***Pseudomonas aeruginosa*** | | | |
| Wild -type PAO1 | 16 | 7 | 5 |
| *ΔF6 cheW* | 15 | 8 | 6 |
| *Δmcp 'mcpA' (PAO180)* | 8 | 4 | 2 |
| *ΔF7 cheW (PAO177)* | 33 | 12 | 0 |
| *ΔF7 cheA (PAO 178)* | 34 | 24 | 0 |
| *Δmcp 'aer2', 'mcpB' (PAO176)* | 21 | 11 | 0 |
| ***Shewanella oneidensis MR-1*** | | | |
| Chemostat growth | 29 | 20 | 7 |
| Batch culture growth | 29 | 18 | 0 |
| ***Methylomicrobium alcaliphilum 20Z*** | | | |
| Wild-type | 8 | 5 | 2 |

**Table S2.**
Chemosensory gene clusters in the genomes of *V. cholerae*, *P. aeruginosa*, *S. oneidensis* and *M. alcaliphilum*.

| | Classification | Alternative name in literature | Function | Gene cluster |
|---|---|---|---|---|
| ***Vibrio cholerae*** | | | | |
| Cluster I | F9 | - | Unknown | VC1394-VC1405 |
| Cluster II | F6 | - | Chemotaxis | VC2059-VC2064 |
| Cluster III | F7 | - | Unknown | VCA1090-VCA1095 |
| ***Pseudomonas aeruginosa*** | | | | |
| Cluster I/V | F6 | Che I | Chemotaxis | PA1457-PA1464 |
| Cluster II | F7 | Che II | Unknown | PA0173-PA0180 |
| Cluster III | ACF | Wsp | Biofilm formation | PA3703-PA3708 |
| Cluster IV | TFP | Chp | Twitching motility | PA0410-PA0415 |
| ***Shewanella oneidensis*** | | | | |
| Cluster I | F7 | CheA-1 | Unknown | SO_2117-SO_2126 |
| Cluster II | F6 | CheA-3 | Chemotaxis | SO_3200-SO_3209 |
| ***Methylomicrobium alcaliphilum*** | | | | |
| Cluster I | F7 | - | Unknown | MEALZ_2869-MEALZ_2879 |
| Cluster II | F8 | - | Unknown | MEALZ_2939 - MEALZ_2942 |
| Cluster III | F6 | - | Unknown | MEALZ_3148 - MEALZ_3158 |

**Table S3.**

Locations of electron density layers in arrays. Distances are measured from the CheA/CheW baseplate in nanometers. Uncertainties reported are the expanded standard uncertainty.hemosensory gene clusters in the genomes of *V. cholerae*, *P. aeruginosa*, *S. oneidensis* and *M. alcaliphilum*.

| **Layer** | ***V. cholerae*** | ***P. aeruginosa*** | ***S. oneidensis*** | ***M. alcaliphilum*** |
|---|---|---|---|---|
| Inner membrane (IM) | 38.4±1.9 nm | 40.3±1.8 nm | 35.5±2.7 nm | 35.1±2.8 nm |
| Layer 3 (L3) | 29.5 ±1.9 nm | 30.7±1.8 nm | 30.7±4.3 nm | - |
| Layer 2 (L2) | 24.8±1.9 nm | 24.0±1.8 nm | 24.0±2.7 nm | 25.3±4.4 nm |
| Layer 1 (L1) | 17.5±1.9 nm | 17.9±2.9 nm | 17.3±2.7 nm | 17.4±4.4 nm |
| Signaling Layer (SL) | - | 7.0±1.8 nm | 6.7±4.3 nm | 7.5±2.8 nm |

**Table S4.**

*P. aeruginosa* strains

| Strain name | PA ORF | Gene |
|---|---|---|
| PW1307 | PA0178 | CheA F7 system |
| PW1305 | PA0177 | CheW F7 system |
| PW1312 | PA0180 | MCPA |
| PW3654 | PA1464 | CheW F6 system |

**Table S5.**

Cryo-Electron Tomograms used in this study available on ETDB. Tomograms can be found on ETDB by the Open Index Protocol id (OIP id).

| Jensen Lab id | OIP id | Organism |
|---|---|---|
| ab2015-06-02-1 | 6cef7c25e66ce42e4e2440490a68845a551303<br>77f7bc9e096c48fd83b47ae9cd | *Vibrio cholerae* |
| ab2015-06-02-2 | 98d93b8c8e390eb6c200fc2d30c9626cb1a7c1<br>40cd930a60eddb3a179aa8226b | *Vibrio cholerae* |
| ab2015-06-02-3 | 39a5d8490ee3823eaf785268aba1593d3be05b<br>8bf6cc029d2e0c247bb7b0a6ba | *Vibrio cholerae* |
| ab2015-06-02-5 | 2620f1ee27c5e63853bb3f3d09796b19d7bd91<br>fbdb5781daa892d68fb31be735 | *Vibrio cholerae* |
| ab2015-06-02-6 | 9ba344e00d31084d2aef8a799f3089e46ead3b<br>28a02c0e73c5852337f67121fc | *Vibrio cholerae* |
| ab2015-06-02-7 | 1e00525c41a9faaa849befd686b70757a6f25a9<br>ceacbcde951b6eeef99006f09 | *Vibrio cholerae* |
| ab2015-06-02-8 | 3cb8dc276fb373a99af2ee54301fcb0989eb003<br>7b9c3c2bd21f8e035f0f98215 | *Vibrio cholerae* |
| ab2015-06-02-9 | 5739b1525b6a0e906de41a8e024e0b502f964<br>e19193feed1a62ced0e41b5dc17 | *Vibrio cholerae* |
| ab2015-06-02-10 | 7e869e97212b3df06af70decf577ef3836c1ec4<br>787f1cbe677da60aca3bda727 | *Vibrio cholerae* |
| ab2015-06-02-11 | 8329b3a0b5373c89d5280c42ca8aabb19a38a6<br>ee03e5910275b82776ff8745fa | *Vibrio cholerae* |
| ab2015-06-02-12 | de04da0f550b2f37d4a0847e13b637877ed699<br>4bb5d8d0c24c87ad9f3cf52924 | *Vibrio cholerae* |
| ab2015-06-02-13 | 4caf0202a48c92372ca3a24101d9c5e2b446fad<br>bf80109b5d0710598a11fa3aa | *Vibrio cholerae* |
| ab2015-06-02-14 | b2f7beb0de38a1eb8d0eab4d8493616c912293<br>1ec0490e1bb5b45474a032098f | *Vibrio cholerae* |
| ab2015-06-02-15 | bd8198a02bd66929811014f1b15a243f47054<br>33876cb56c784bdc8fb91378c00 | *Vibrio cholerae* |
| ab2015-06-02-16 | 87c298d70069315af599618c344d80bf2fc48f4<br>4c0057002798241842b58b5c9 | *Vibrio cholerae* |
| ab2015-06-02-17 | 066cf7fae820b463b2e7405a9078e97b6de991<br>0587458dfd904759d75fb4ed2c | *Vibrio cholerae* |

| Jensen Lab id | OIP id | Organism |
|---|---|---|
| ab2015-06-02-18 | 82fdc6778793542fad9bfe7c18a6fa972dc3bf23 0b2e108aa53a5807811bdad7 | *Vibrio cholerae* |
| ab2015-06-02-20 | 2524250ba11140c386c211aa14d164a166926 ee9ca7af5566eedbbba29856c48 | *Vibrio cholerae* |
| ab2015-06-02-21 | 95a8202f59bdcfc9814f417aa40b3c355fc6f12c b147784d1d1665d8ccce8221 | *Vibrio cholerae* |
| ab2015-06-02-22 | 09443b1917375d0f2755c3f4c2029dc847c654 efdb2e5a3df0c5938a9afaa1a9 | *Vibrio cholerae* |
| ab2015-06-02-23 | 453536f21d3074fd3f6fcae6d834c6e37335f46 6611e2f1842c6beb8ee77681e | *Vibrio cholerae* |
| ab2015-06-02-24 | 03ab91b142bf62ed9c21e1da4b17e25d7c73b7 0cbd6a25481342d9736f2a2290 | *Vibrio cholerae* |
| ab2015-06-02-25 | 8853aea7df6d8a168b62b9db8694ad82dfa23c 9b00eafb3f71b025f20cb34abd | *Vibrio cholerae* |
| ab2015-06-02-26 | 8770b212ceddc32db6f1ae0b8a40a5eaa1ec42c 31bceda0c4919b86dcde7246b | *Vibrio cholerae* |
| ab2015-06-02-27 | e1b34b3df7aa6367398010bf8b96d5a003759a 2dd71a473e45a43d598e779d29 | *Vibrio cholerae* |
| ab2015-06-02-29 | 08324bf01302b1db927045548218ff3e317aea e74602363e669a187c52e234be | *Vibrio cholerae* |
| ab2015-06-02-30 | 2eda58ca01240b7486bda36c36e36217918c8 9fcb1361ddda13eeda95afc4271 | *Vibrio cholerae* |
| ab2015-06-02-31 | 6b6b8ba7f2127c66614756b85d4bb0efde943c bfc305e40a3330181009bd893e | *Vibrio cholerae* |
| ab2015-06-02-32 | 61c86c9a912eb1bcada440939a48d46e99128f feca99578b16e00e22031f4a22 | *Vibrio cholerae* |
| ab2015-06-03-1 | 197f9f4bbc9f5fa49c4644f3f40df46cbf2b82399 e8137db7799b9e030bc8455 | *Vibrio cholerae* |
| ab2015-06-03-2 | 544970e8e49ff409d4f61730c89757fdbee6fff3 64c1c604d355bc522c8d0f12 | *Vibrio cholerae* |
| ab2015-06-03-3 | e87e0b6e154e5dfec0120ce92e6400764ae249 64aa592553e49de6187b7ffb57 | *Vibrio cholerae* |
| ab2015-06-03-4 | 20d581e5578e3156ddf96496119c1296494e0 62deecf45634c79f19f331368b6 | *Vibrio cholerae* |
| ab2015-06-03-5 | 10c8a0be7b87b3d95eb2d8573e782a856e493 b65735aab290c0a772092d2ecfc | *Vibrio cholerae* |

| Jensen Lab id | OIP id | Organism |
|---|---|---|
| ab2015-06-03-6 | 0c1b01aa1ed45fcba7d17500c4829f9480924fa210f9706d6a1a5da9d4dc1477 | *Vibrio cholerae* |
| ab2015-06-03-7 | 21482339f240527367c699214ba438d2b9a7ac6376d611f2d69e0f4c1d1d0039 | *Vibrio cholerae* |
| ab2015-06-03-8 | 453a055bebf6eb8d65f2154f51a568cfc3f58c36aa445ff6be7a820c0d3963e9 | *Vibrio cholerae* |
| ab2015-06-03-9 | faed3ad567cb4667d1986bb9a7869a4a9da0a9e2d016dbb6d823e15f5d1cc149 | *Vibrio cholerae* |
| ab2015-06-03-10 | 1c8abd7e25085cccce2ce38fb25ff51ac70ddfc88cad18322406467298bfaa3e | *Vibrio cholerae* |
| ab2015-06-03-11 | 920321536f97c55c300ad77d279265973442bd4ec0e7286a95f2d134ec384b42 | *Vibrio cholerae* |
| ab2015-06-03-12 | 9262dbfb593f79bf0c9cac6f9125326ede755dd3e4edc569b68b0fc279f84d8d | *Vibrio cholerae* |
| ab2015-06-03-13 | 72b627c7010d0ea37a8aa011925069437195c51f77e5adcf41f6a8330e75cc65 | *Vibrio cholerae* |
| ab2015-06-03-14 | 223d692a658a507a7085c75f8c757b05c1cf1d2ee8e352a1db4371f2062e3ec4 | *Vibrio cholerae* |
| ab2015-06-03-15 | c55cf52f955779f23cda1c2244e2b64315f8fbb490f10689f9c32b7a87a99512 | *Vibrio cholerae* |
| ab2015-06-03-16 | b59e667fadb1914c70ad191dad31a30a87ba416e8de4877000d1f6cb52869601 | *Vibrio cholerae* |
| ab2015-06-03-17 | 067ac5b8ce49dffe5254c958a0ef6bfd601de8778c47f17d8db6c76f2b03d882 | *Vibrio cholerae* |
| ab2015-06-03-18 | c9dd866db34f7ed34af143d2a879e5b28035fce2a4392aceb54f7824ceec0e37 | *Vibrio cholerae* |
| ab2015-06-03-19 | fabd724070997b695344be1bc4ad91faa2b2c82876f6bda65aeeee18eb39ca29 | *Vibrio cholerae* |
| ab2015-06-03-20 | 34ff16d0af7b7a6cfbe49d20981049070d2b9b9f5a1919896a839eb62dc8c6a8 | *Vibrio cholerae* |
| ab2015-06-03-21 | 4440060c9b511b30ec0b7c123b3891a3823a77a8393437862aa5344392e0be26 | *Vibrio cholerae* |
| ab2015-06-03-22 | 48c976b5ec236342fdd4aefe521086e3657da4a45c4afad6003efdb44a5e8275 | *Vibrio cholerae* |
| ab2015-06-03-23 | 1c1e69acb1c51ec5655f2fc7e1296de6494c1c68785e7f4a7ce0cc651d2125c9 | *Vibrio cholerae* |

| Jensen Lab id | OIP id | Organism |
|---|---|---|
| ab2015-06-03-24 | aa070936e1aee30b373bc2d2aed88f1576cd3e<br>d6766dbc9f92f120be46aeeabe | *Vibrio cholerae* |
| ab2015-06-03-25 | d84b38caf03c63be65666a628fa8f90aeaec899<br>1ba29dad0b539585fbcf3b96c | *Vibrio cholerae* |
| ab2015-06-03-26 | f38ac8c272b532e7dd3e3da38b1fff38f42385a<br>2aa55872dda24bf10720544ba | *Vibrio cholerae* |
| ab2015-06-03-27 | 9d8070fbfbab0e5bfdd555544219789104fb5d<br>cbda613d24385c416ccd3c3158 | *Vibrio cholerae* |
| ab2015-06-03-28 | bd0aaf3c9bb4904c623c2304b5ca71f3e7dbbb<br>d15fdbe2eb481dbdb01d561986 | *Vibrio cholerae* |
| ab2015-06-03-29 | 81345effbcb7600a9f57a2f556a10f3a20a6845<br>df3ca51ccc1d67d964ee8e21f | *Vibrio cholerae* |
| ab2015-06-04-5 | e95c286a1ca5b5b3b3b5bf0a318a30c627623d<br>2b63b68c9991dab337ae48e8c2 | *Vibrio cholerae* |
| ab2015-06-04-6 | 79b0258243f18a1752dac35aa7e0bff1a711f79<br>1ddd0379c184e05d9d1c5dd86 | *Vibrio cholerae* |
| ab2015-06-04-7 | 6b8f4ae3d17ab82176fe1bb2c79c55d5ce757c<br>5355323bbadc84d17a5a6f75dd | *Vibrio cholerae* |
| ab2015-06-04-8 | 18fca4c49057b64129c9dbd0f9c8be7562fcee4<br>d5fff21a449b1fd79a5e910f8 | *Vibrio cholerae* |
| ab2015-06-04-9 | a9e50ed3ce23097a99fdaa4d949f5353bcba9f5<br>ce3cd7a49d5f3df08d668a989 | *Vibrio cholerae* |
| ab2015-06-04-10 | 275fbd79f1fb013309e3dc4a22c607ab406b6b<br>2a16489a508c3490ede71dac93 | *Vibrio cholerae* |
| ab2015-06-04-11 | eadc30e57a5bbceba8e69544d43debdb4d7139<br>2ef20f16c8d9a5659a2dc877a9 | *Vibrio cholerae* |
| ab2015-06-04-12 | 82b9550d3ebda9a078ece5cea3cfd02ceece06d<br>f9115d8b2d558d5b5c073f9dd | *Vibrio cholerae* |
| ab2015-06-04-13 | 204dea6006208b691ae45b1aa8a62421894c0<br>d956a5d5bf6aaa8a3e75253345d | *Vibrio cholerae* |
| ab2015-06-04-14 | 7504bd89ddb0fd83ed926065c9d2cf7b57db00<br>72ebb740cfefdc957f1d5c4c81 | *Vibrio cholerae* |
| ab2015-06-04-15 | 7bc4afbf390105748d85295198579497fd0d67<br>12da0eb190eec2436cf6617349 | *Vibrio cholerae* |
| ab2015-06-04-16 | c99f6e5934554670fe2166f2f8deb7ad17f4536<br>96c914b4ca1ea88f7e009f0a4 | *Vibrio cholerae* |

| Jensen Lab id | OIP id | Organism |
|---|---|---|
| ab2015-06-04-17 | 095901997ad173f1a9cf5804cd7f92e2c95ee7b d4be14b1fa1fe94ed74e6a85d | *Vibrio cholerae* |
| ab2015-06-04-18 | 2f236dac0a263b2e3278aef8ab2aad6c7ace83b de857c6916683146369424e93 | *Vibrio cholerae* |
| ab2015-06-04-19 | b4daca6e8f77b714d01faac2b7dc9933c4617d 736b0a6ef29eb9e1056c0214f8 | *Vibrio cholerae* |
| ab2015-06-04-20 | ed95dd31b7e3a97a2bedc0e8985ad471dad7c7 447a7070d5d5b37b8f259da41a | *Vibrio cholerae* |
| ab2015-06-04-21 | a41257cc5f1d3c4a84cd9645921671bff4e5b58 de593872740a35172fd4102a5 | *Vibrio cholerae* |
| ab2015-06-04-22 | 863a029fcd72521b29fa741b7a13b2df4420fcc 9491a090eab709c7c7d727514 | *Vibrio cholerae* |
| ab2015-06-04-23 | c63edc39ab954f54d3ec80607005b54739f694 0f8e317c3ea88d951b0e63ed82 | *Vibrio cholerae* |
| ab2015-06-04-24 | 5f040233e108f1d0d81448d4b7c44476947e2 075742fd79ad2f0d0adf9f686fa | *Vibrio cholerae* |
| ab2015-06-04-25 | 9a31271d398507a3e7cb0694927dbda47043f 35bf8171cbd4f56df5896c7e060 | *Vibrio cholerae* |
| ab2015-06-04-26 | 687d83b774894a40b9f35c54e0e50d2e11613 d6260723ab83573e3fd6c55b309 | *Vibrio cholerae* |
| ab2015-06-04-27 | 5cc452dfbe2e22d09dd2102b010e926f7c9735 5c0c209148f049292cf3b910de | *Vibrio cholerae* |
| ab2015-06-04-28 | 49ffb9f6a24a7d07799479b844a01ed91a204f9 5afa28a8d59561b858d9a4d19 | *Vibrio cholerae* |
| ab2015-06-04-29 | 38f8ed8a6cba9fdab947246fd283023b21353f1 66c6e41f949320a028cea2f1a | *Vibrio cholerae* |
| ab2015-06-04-30 | 92c05f4866a31ee526bac1d95d513c9d0063e7 94554cdc124651fabc1e8c9395 | *Vibrio cholerae* |
| ab2015-06-04-31 | 60d87f38e11916f13cf6345b22f0b5fc567d6ec ba651e7ed98c1ba23faca1a27 | *Vibrio cholerae* |
| ab2015-06-04-32 | 2b200698dc605f2234c28d33d12a2eeed4bfb6 808399563791bf228fda34936e | *Vibrio cholerae* |
| ab2015-05-29-38 | 71b7016135114ee9988faf24a00aec627349d8 610625a8925152e418ce042cb8 | *Pseudomonas aeruginosa* |
| ab2015-05-29-33 | 3bc9419d595c59d2ffb177294ceb038f5c4a402 87cc1d2ff95ab43e42e22c5b5 | *Pseudomonas aeruginosa* |

| Jensen Lab id | OIP id | Organism |
|---|---|---|
| ab2015-05-29-34 | baf59323b4cd6b20e471b0587f4349bb437cc8 9feba37eb726c76d2b35996859 | *Pseudomonas aeruginosa* |
| ab2015-05-29-35 | 375ec0e6c3542bc3d33e7ec94c751bd1669ffe9 984a4ccb8948ddc246bd02517 | *Pseudomonas aeruginosa* |
| ab2015-05-29-36 | dda73882e77f51644ba4d9fc73810f58c0789cf 5ebd5c4c64108940c978d3913 | *Pseudomonas aeruginosa* |
| ab2015-05-29-37 | e0d1b4daa89125bfae9539157ab0d459e9a10b 53148d17af60eb3afadf667e37 | *Pseudomonas aeruginosa* |
| ab2015-05-29-38 | 71b7016135114ee9988faf24a00aec627349d8 610625a8925152e418ce042cb8 | *Pseudomonas aeruginosa* |
| ab2015-05-29-39 | 9ff1533aa7d28d3d64439293ea263ca3ce6a61 01f19c7367426d9a3bba3773e6 | *Pseudomonas aeruginosa* |
| ab2015-05-29-40 | c5f0fcf46599f6ee48fe23a59fc1d752cd64ab40 31c3846ffb85d078a8bfaf5c | *Pseudomonas aeruginosa* |
| ab2015-05-29-41 | fe52314687302e52d76a76292a84a8e7730dac 37451c7026621190f2185cd7df | *Pseudomonas aeruginosa* |
| ab2015-05-29-42 | fc8e9658e5fe5d5e30364f389fc123f474537bd a5132fb71b12af99aaf02e7a2 | *Pseudomonas aeruginosa* |
| ab2015-05-29-43 | 433cf08b0ab2269a91f4e946b17c551e8a92a3 0c8c5214bc8384072fc8915b85 | *Pseudomonas aeruginosa* |
| ab2015-05-29-44 | 9162280a2e505e7a74c03de6202f084eb4b838 82453525539fdf87da3905d77c | *Pseudomonas aeruginosa* |
| ab2015-05-29-45 | 9db0388f6a5482eb6cad84cd7abf0ddad7bcd4 a3d741d971cc9dd3a2c378cda2 | *Pseudomonas aeruginosa* |
| ab2015-05-29-46 | 31eee1b22c3f3010a3a31df973be75ff1a12606 9e78bfbb1dbed30cd3985330d | *Pseudomonas aeruginosa* |
| ab2015-05-29-47 | 799edd343b37e6dff51118d39052ad7241e7c0 5238286bcc779c1ab9227f8a4c | *Pseudomonas aeruginosa* |
| ab2015-05-29-48 | 72025551790f3ada513cffe56e1f0971b3844f1 c985e4f3e3b7579dd349aecd1 | *Pseudomonas aeruginosa* |
| ab2015-05-29-49 | 76b1c8f739812662c22aee910f2e1a2bc7ac72f 9c79ee19df0493c60be7f6711 | *Pseudomonas aeruginosa* |
| ab2015-05-29-50 | 2f3db7b47e7d42594be348214e07578d7ea1f0 8a10f8cd937bec3abb8692c6aa | *Pseudomonas aeruginosa* |
| ab2015-05-29-51 | dbe07cf9023a7c8d67cea0f6bc02eeaea93f5c2e 60453ca2aa4685d8e3a53654 | *Pseudomonas aeruginosa* |

| Jensen Lab id | OIP id | Organism |
|---|---|---|
| ab2015-05-29-52 | 8f861e2f49e3b2155f99422428acb7ca764c5c2 2352dd6f5c411d89daaa99c8c | *Pseudomonas aeruginosa* |
| ab2015-05-29-53 | 8bbc0699df1ecaeff06813ea816ded484d90867 9412e938d42afbb30e0f47747 | *Pseudomonas aeruginosa* |
| ab2015-05-29-54 | 87be5d50a167d3c23ad2126bc4d72844c0ed6 af2848768fd1bb764ad0a05355b | *Pseudomonas aeruginosa* |
| ab2015-05-29-55 | 975a50a6ac4a0a1ce191d3f339c433b73ba15a 3a8eff22b3768a9327e1ca5a21 | *Pseudomonas aeruginosa* |
| ab2015-05-29-56 | d0799bdb371912d4370d6226ae8263e1f9bf4 19c49c42c16514c657f11f05f37 | *Pseudomonas aeruginosa* |
| ab2015-05-29-57 | dfd11ff2b7b5261968452f994235bece1d208b 793f0c7768b883163fa0cad72c | *Pseudomonas aeruginosa* |
| ab2015-05-29-58 | 06dc2aa14271a7d5c79c6e112a3aeacc8c6432 46c659fab33b3bbb01fbe25356 | *Pseudomonas aeruginosa* |
| ab2015-05-29-59 | f92bcceb78dac999467e57653d9e3ab4017261 8fc09fb5806e64040e8a3410f5 | *Pseudomonas aeruginosa* |
| ab2015-05-29-60 | bd0b6b93ba4f3528fe3f78a789dd847ee83a6d 4b08cd6e652053e72aefa6897b | *Pseudomonas aeruginosa* |
| ab2015-05-29-61 | 37d4d1c6d5229a5db3700596859b305aea3aa 460d67362ba176424c9217ebf28 | *Pseudomonas aeruginosa* |
| ab2015-05-29-62 | 4e059367636fa821409e894b49151556c01d6 5ebe01eca3e4d4fedec6531adc7 | *Pseudomonas aeruginosa* |
| ab2014-010-14-1 | b9d6746d36ee7943e0eb19c9299214e60080c 78b367558d6748820c84c2aec76 | *Pseudomonas aeruginosa* |
| ab2014-010-14-10 | c8c1989c09901dd7a257d86497a713e85a761 edd2209efe0d8784d228e5b3585 | *Pseudomonas aeruginosa* |
| ab2014-010-14-11 | 18ec447aac947d6ed610084fbb9e808bdb761b f68919b853ac4946632d6c6e50 | *Pseudomonas aeruginosa* |
| ab2014-010-14-12 | 58c0a2aab1b8f0322fcc4b93838458a820fc8db 74f716b53ba166ae2cbe92001 | *Pseudomonas aeruginosa* |
| ab2014-010-14-13 | 9e8108ef3cf66406b628ec973b10079efc0ca57 81a05ca1a875ed6fdecc2fe02 | *Pseudomonas aeruginosa* |
| ab2014-010-14-14 | 7935084f249ec5bc222b38951cd369dab76cfa b95a769d42d0eb1312cc5bb50c | *Pseudomonas aeruginosa* |
| ab2014-010-14-15 | 51b27c3cf74ce394c1d54773bd1a7d15d3379f c8db4aa2f8b9dbdbaabdd71251 | *Pseudomonas aeruginosa* |

| Jensen Lab id | OIP id | Organism |
|---|---|---|
| ab2014-010-14-17 | 764f7220c7323438d65b7d4334916e9aadb0b e04be1b662615be1ad1312407c6 | *Pseudomonas aeruginosa* |
| ab2014-010-14-22 | 797a818b0722e811f5ed0d6d22bdfdb158fabb cd22e8bfc7f1010ad265af8897 | *Pseudomonas aeruginosa* |
| ab2014-010-14-23 | 322a5ff8d6dda5854aaafec4883226c8463c2e9 a08ce553c969ca6b98cfedef9 | *Pseudomonas aeruginosa* |
| ab2014-010-14-3 | 2c11ded42bbc0cd87dc67464474187c3c82d7 1742a8b50f346516a958361706a | *Pseudomonas aeruginosa* |
| ab2014-010-14-4 | 20cd19c0fa4b3e32b33075348be5ab3102def5 083cce6fd7ac55b01ac0cb5835 | *Pseudomonas aeruginosa* |
| ab2014-010-14-5 | 3e6133278fc31ba361c1cd7c95c081a822d972 3dc3b7f3549870510a422e2522 | *Pseudomonas aeruginosa* |
| ab2014-010-14-6 | 3cbeac672c18cd66252b47d65f0ec124b17baf5 19849a6844fab4af70fbbcea1 | *Pseudomonas aeruginosa* |
| ab2014-010-14-8 | cb237b8021a9e653613163394c11258bb3b72 fe09720a977596c18478e5d43c2 | *Pseudomonas aeruginosa* |
| ab2014-010-14-9 | 8bf9ec4e43a3e6bd11c44bb6d341d138e14b41 c776e66e21a8a17d399c1c69c2 | *Pseudomonas aeruginosa* |
| sc2014-02-14-1 | ecfe996b6c698beb104f0cf84bda7955a48c7ed b6049ce2b9256cb3b1e2b85b0 | *Methylomicrobium alcaliphylum* |
| sc2014-02-12-1 | 37a38f6896b2fbc3379ebba9976e69ccefd45da a011b009f9fc498f40ab5b68e | *Methylomicrobium alcaliphylum* |
| sc2014-02-12-3 | 82c7e7111fad00bd9de9be253c966a0d87b3b5 85cf47a4e3ada52b7a90ae362f | *Methylomicrobium alcaliphylum* |
| sc2014-02-12-6 | c807ab328c5bd06874ad4f9fb731fafd888f238 94d00cafed1089f988407ed99 | *Methylomicrobium alcaliphylum* |
| sc2014-02-12-7 | 99a62dfe3b6bedf92f6bd93916b1fe055c1abd7 d4d2f781348c01ae89c45dbd5 | *Methylomicrobium alcaliphylum* |
| sc2014-02-12-8 | 6ce0a323a040374567356e08c438fd42806aef 88c73f105e2d9e13684d940ae8 | *Methylomicrobium alcaliphylum* |
| sc2014-02-12-9 | 9e828a83fb3538f1f6f9942ea4216cd7fe20ebcc 1c82ed3dba8f7e34bdc3a1e9 | *Methylomicrobium alcaliphylum* |
| sc2014-02-12-12 | 262b60a13fbcd486cc7c4ac73b739ddccb23d9 3cf8d5d4138ae911af40f95480 | *Methylomicrobium alcaliphylum* |

**Table S6.**
Atomic models used to produce the homology models used in this work.

| Domain(s) | PDB code | Reference |
|---|---|---|
| HAMP(*A. fulgidus*) + MCP_Signal(*E. coli*) | 3ZX6 | (*74*) |
| PAS(*P. aeruginosa*) | 4HI4 | (*22*) |
| PAS + HAMP (*P. aeruginosa*) | 3VOL | (*75*) |
| 3xHAMP (*P. aeruginosa*) | 4I3M | (*76*) |

**Table S7.**

Relevant files used to build the homology models produced in this work.

| File | Type | Description |
|---|---|---|
| 3XZ6_4I3M.pir | Sequence alignment | Sequence alignment used to build the 2H+S homology model |
| 3XZ6_4I3M_74.pdb | 3D atomic model | Best homology model from 2H+S |
| 4HI4_BD.pdb | 3D atomic model | chains B and D of 4HI4 aligned with 2H+S model |
| 3ZX6_4I3M_4HIH.pir | Sequence alignment | Sequence alignment used to build the P+2H+S homology model |
| 3ZX6_4I3M_4HIH_99.pdb | 3D atomic model | Best homology model from P+2H+S |
| 4I3M.bio.pos.pdb | 3D atomic model | Model of 4I3M positioned against 3ZX6_4I3M_4HIH_99.pdb to build the model for Aer2 (PA0176) |
| 3ZX6_4I3M_4HI4_4I3M.pir | Sequence alignment | Sequence alignment used to build the model for Aer2 (PA0176) |
| Aer2Pa_3HAMP_PAS_2HAMP.B99990041.pdb | 3D atomic model | Best Aer2 (PA0176) homology model |
| 3ZX6_4I3M_4HI4_4HI4.pir | Sequence alignment | Sequence alignment used to build the model for Aer2-like (VCA1092) |
| VCA1092.B999900035.pdb | 3D atomic model | Best Aer2-like (VCA1092) homology model |
| 3ZX6_4I3M_4HI4_4HI4_SO.pir | Sequence alignment used to build the model for Aer2-like (SO_2123) | |

| File | Type | Description |
|---|---|---|
| SO_2123.B99990017.pdb | 3D atomic model | Best Aer2-like (SO_2123) homology model |
| hamp_sequence_for_MEALZ.linsi.fa | Sequence alignment | Sequence alignment of HAMP domains in the group of Pseudomonas group similar to the 3 HAMPs in 4I3M and the C-terminal HAMP of MEALZ_2872 |
| RAxML_bipartitions_50coll.hamp_sequence_for_MEALZ.linsi.rec.tree | Phylogenetic Tree | The tree with maximum likelihood based on hamp_sequence_for_MEALZ.linsi.fa |
| 4I3M.bio.HAMP2.withtail4alignment.pdb | 3D atomic model | Model of the second HAMP of 4I3M with part of the helix connecting to the third HAMP. |
| 4I3M.bio.HAMP2.alnMEALZ.pdb | 2\3D atomic model | Model of the second HAMP of 4I3M without part of the helix connecting to the third HAMP. |
| 3ZX6_4I3M_4HI4_HAMP2_MEALZ.pir | Sequence alignment used to build the model for Aer2-like (MEALZ_2872) | |
| MEALZ_2872_wHAMP.B99990020.pdb | 3D atomic model | Best Aer2-like (MEALZ_2872) homology model |

**Table S8.**

310 randomly selected non-redundant γ-Proteobacteria genomes used in this work. The presence of an F7 system is indicated.

| **Genome** | **has F7** |
|---|---|
| *Acidithiobacillus caldus SM-1* | no |
| *Acidithiobacillus ferrivorans SS3* | no |
| *Acidithiobacillus sp. GGI-221* | no |
| *Acidithiobacillus thiooxidans ATCC 19377* | no |
| *Acinetobacter baumannii AB5075* | no |
| *Acinetobacter bereziniae LMG 1003* | no |
| *Acinetobacter calcoaceticus RUH2202* | no |
| *Acinetobacter haemolyticus ATCC 19194* | no |
| *Acinetobacter johnsonii SH046* | no |
| *Acinetobacter junii SH205* | no |
| *Acinetobacter lwoffii SH145* | no |
| *Acinetobacter nosocomialis Ab22222* | no |
| *Acinetobacter oleivorans DR1* | no |
| *Acinetobacter parvus DSM 16617 = CIP 108168* | no |
| *Acinetobacter radioresistens DSM 6976 = NBRC 102413* | no |
| *Acinetobacter sp. NCTC 10304* | no |
| *Acinetobacter ursingii DSM 16037 = CIP 107286* | no |
| *Aeromonas aquariorum AAK1* | no |
| *Aeromonas caviae Ae398* | no |
| *Aeromonas hydrophila SSU* | no |
| *Aeromonas media WS* | no |
| *Aeromonas salmonicida subsp. salmonicida A449* | no |
| *Aeromonas veronii AER397* | no |
| *Alcanivorax borkumensis SK2* | no |

| **Genome** | **has F7** |
|---|---|
| *Alcanivorax dieselolei B5* | yes |
| *Alcanivorax hongdengensis A-11-3* | no |
| *Alcanivorax pacificus W11-5* | yes |
| *Alcanivorax sp. DG881* | no |
| *Aliivibrio salmonicida LFI1238* | no |
| *Alishewanella aestuarii B11* | no |
| *Alishewanella agri BL06* | no |
| *Alishewanella jeotgali KCTC 22429* | no |
| *Alkalilimnicola ehrlichii MLHE-1* | no |
| *Allochromatium vinosum DSM 180* | yes |
| *Alteromonadales bacterium TW-7* | yes |
| *Alteromonas mediterranea MED64* | no |
| *Alteromonas sp. SN2* | yes |
| *Azotobacter vinelandii DJ* | yes |
| *Beggiatoa alba B18LD* | yes |
| *Beggiatoa sp. SS* | yes |
| *Cardiobacterium hominis ATCC 15826* | no |
| *Cardiobacterium valvarum F0432* | no |
| *Cellvibrio japonicus Ueda107* | no |
| *Cellvibrio sp. BR* | yes |
| *Chromohalobacter salexigens DSM 3043* | yes |
| *Citrobacter freundii 4_7_47CFAA* | yes |
| *Citrobacter koseri ATCC BAA-895* | yes |
| *Citrobacter rodentium ICC168* | yes |
| *Citrobacter sp. 30_2* | yes |
| *Citrobacter youngae ATCC 29220* | yes |

| **Genome** | **has F7** |
|---|---|
| *Colwellia psychrerythraea 34H* | no |
| *Cronobacter sakazakii ES15* | yes |
| *Cronobacter turicensis z3032* | yes |
| *Dichelobacter nodosus VCS1703A* | no |
| *Dickeya dadantii Ech703* | yes |
| *Dickeya zeae Ech1591* | yes |
| *Ectothiorhodospira sp. PHS-1* | yes |
| *Edwardsiella ictaluri 93-146* | yes |
| *Edwardsiella tarda ATCC 23685* | yes |
| *Endoriftia persephone 'Hot96_1+Hot96_2'* | no |
| *Enhydrobacter aerosaccus SK60* | no |
| *Enterobacter asburiae LF7a* | yes |
| *Enterobacter cancerogenus ATCC 35316* | yes |
| *Enterobacter cloacae subsp. cloacae GS1* | yes |
| *Enterobacter hormaechei ATCC 49162* | yes |
| *Enterobacter radicincitans DSM 16656* | yes |
| *Enterobacter sp. 638* | yes |
| *Enterobacteriaceae bacterium 9_2_54FAA* | yes |
| *Erwinia amylovora CFBP1430* | yes |
| *Erwinia billingiae Eb661* | yes |
| *Erwinia pyrifoliae Ep1/96* | yes |
| *Erwinia sp. Ejp617* | yes |
| *Erwinia tasmaniensis Et1/99* | yes |
| *Escherichia albertii TW11588* | yes |
| *Escherichia coli KTE229* | yes |
| *Escherichia fergusonii ECD227* | yes |

| Genome | has F7 |
|---|---|
| *Escherichia hermannii NBRC 105704* | yes |
| *Escherichia sp. TW09276* | yes |
| *Ferrimonas balearica DSM 9799* | no |
| *Fluoribacter dumoffii Tex-KL* | no |
| *Frateuria aurantia DSM 6220* | yes |
| *Gallaecimonas xiamenensis 3-C-1* | yes |
| *Glaciecola agarilytica NO2* | yes |
| *Glaciecola arctica BSs20135* | no |
| *Glaciecola chathamensis S18K6* | yes |
| *Glaciecola lipolytica E3* | no |
| *Glaciecola mesophila KMM 241* | no |
| *Glaciecola nitratireducens FR1064* | yes |
| *Glaciecola pallidula DSM 14239 = ACAM 615* | yes |
| *Glaciecola polaris LMG 21857* | no |
| *Glaciecola psychrophila 170* | no |
| *Glaciecola sp. 4H-3-7+YE-5* | yes |
| *Grimontia hollisae CIP 101886* | yes |
| *Grimontia sp. AK16* | yes |
| *Hafnia alvei ATCC 51873* | yes |
| *Hahella chejuensis KCTC 2396* | yes |
| *Halomonas boliviensis LC1* | yes |
| *Halomonas elongata DSM 2581* | yes |
| *Halomonas sp. GFAJ-1* | yes |
| *Halomonas titanicae BH1* | yes |
| *Halorhodospira halophila SL1* | yes |
| *Halothiobacillus neapolitanus c2* | no |

| Genome | has F7 |
|---|---|
| *Hydrocarboniphaga effusa AP103* | yes |
| *Idiomarina loihiensis L2TR* | no |
| *Idiomarina xiamenensis 10-D-4* | no |
| *Kangiella koreensis DSM 16069* | no |
| *Klebsiella aerogenes KCTC 2190* | yes |
| *Klebsiella pneumoniae subsp. pneumoniae HS11286* | no |
| *Legionella drancourtii LLAP12* | yes |
| *Legionella longbeachae NSW150* | no |
| *Legionella pneumophila subsp. pneumophila* | no |
| *Listonella anguillarum M3* | yes |
| *Marichromatium purpuratum 984* | yes |
| *Marinobacter adhaerens HP15* | no |
| *Marinobacter algicola DG893* | no |
| *Marinobacter hydrocarbonoclasticus ATCC 49840* | no |
| *Marinobacter hydrocarbonoclasticus VT8* | no |
| *Marinobacter manganoxydans MnI7-9* | no |
| *Marinobacter santoriniensis NKSG1* | no |
| *Marinobacter sp. ELB17* | no |
| *Marinobacterium stanieri S30* | yes |
| *Marinomonas mediterranea MMB-1* | yes |
| *Marinomonas posidonica IVIA-Po-181* | no |
| *Marinomonas sp. MWYL1* | yes |
| *Methylobacter tundripaludum SV96* | yes |
| *Methylococcus capsulatus str. Bath* | no |
| *Methylomicrobium album BG8* | yes |
| *Methylomicrobium alcaliphilum 20Z* | yes |

| **Genome** | **has F7** |
| --- | --- |
| *Methylomonas methanica MC09* | yes |
| *Methylophaga aminisulfidivorans MP* | yes |
| *Methylophaga frappieri* | no |
| *Methylophaga lonarensis MPL* | no |
| *Methylophaga thiooxydans DMS010* | no |
| *Moraxella macacae 0408225* | no |
| *Morganella morganii subsp. morganii KT* | yes |
| *Moritella sp. PE36* | yes |
| *Nitrosococcus halophilus Nc 4* | no |
| *Nitrosococcus oceani ATCC 19707* | no |
| *Nitrosococcus watsonii C-113* | no |
| *Oceanimonas sp. GK1* | no |
| *Pantoea agglomerans 299R* | yes |
| *Pantoea ananatis LMG 20103* | yes |
| *Pantoea sp. aB* | yes |
| *Pantoea stewartii subsp. stewartii DC283* | yes |
| *Pantoea vagans C9-1* | yes |
| *Pectobacterium atrosepticum SCRI1043* | yes |
| *Pectobacterium carotovorum subsp. brasiliensis PBR1692* | yes |
| *Pectobacterium sp. SCC3193* | yes |
| *Pectobacterium wasabiae CFBP 3304* | yes |
| *Photobacterium damselae subsp. damselae CIP 102761* | no |
| *Photobacterium leiognathi subsp. mandapamensis svers.1.1.* | no |
| *Photobacterium profundum SS9* | no |
| *Photobacterium sp. AK15* | no |
| *Photorhabdus asymbiotica* | yes |

| Genome | has F7 |
|---|---|
| *Photorhabdus luminescens subsp. laumondii TTO1* | yes |
| *Proteus mirabilis WGLW6* | yes |
| *Proteus penneri ATCC 35198* | yes |
| *Providencia alcalifaciens DSM 30120* | yes |
| *Providencia burhodogranariea DSM 19968* | yes |
| *Providencia rettgeri Dmel1* | no |
| *Providencia rustigianii DSM 4541* | yes |
| *Providencia stuartii ATCC 25827* | yes |
| *Pseudoalteromonas arctica A 37-1-2* | no |
| *Pseudoalteromonas atlantica T6c* | no |
| *Pseudoalteromonas citrea NCIMB 1889* | yes |
| *Pseudoalteromonas haloplanktis ANT/505* | yes |
| *Pseudoalteromonas luteoviolacea B = ATCC 29581* | yes |
| *Pseudoalteromonas marina mano4* | yes |
| *Pseudoalteromonas piscicida JCM 20779* | yes |
| *Pseudoalteromonas rubra ATCC 29570* | yes |
| *Pseudoalteromonas sp. Bsw20308* | yes |
| *Pseudoalteromonas spongiae UST010723-006* | yes |
| *Pseudoalteromonas undina NCIMB 2128* | yes |
| *Pseudomonas aeruginosa LESB58* | yes |
| *Pseudomonas avellanae BPIC 631* | no |
| *Pseudomonas brassicacearum subsp. brassicacearum NFM421* | no |
| *Pseudomonas denitrificans ATCC 13867* | yes |
| *Pseudomonas entomophila L48* | no |
| *Pseudomonas extremaustralis 14-3 substr. 14-3b* | no |
| *Pseudomonas fluorescens F113* | no |

| Genome | has F7 |
|---|---|
| *Pseudomonas fragi A22* | no |
| *Pseudomonas fulva 12-X* | no |
| *Pseudomonas fuscovaginae UPB0736* | no |
| *Pseudomonas geniculata N1* | yes |
| *Pseudomonas mendocina ymp* | no |
| *Pseudomonas monteilii SB3078* | no |
| *Pseudomonas poae RE*1-1-14* | no |
| *Pseudomonas protegens CHA0* | no |
| *Pseudomonas pseudoalcaligenes KF707* | yes |
| *Pseudomonas psychrotolerans L19* | no |
| *Pseudomonas putida GB-1* | no |
| *Pseudomonas resinovorans NBRC 106553* | yes |
| *Pseudomonas sp. TKP* | no |
| *Pseudomonas stutzeri KOS6* | no |
| *Pseudomonas syringae pv. phaseolicola 1448A* | no |
| *Pseudomonas viridiflava UASWS0038* | no |
| *Pseudoxanthomonas spadix BD-a59* | no |
| *Pseudoxanthomonas suwonensis 11-1* | yes |
| *Psychrobacter arcticus 273-4* | no |
| *Psychrobacter cryohalolentis K5* | no |
| *Psychrobacter sp. PRwf-1* | no |
| *Psychromonas sp. CNPT3* | no |
| *Rahnella aquatilis CIP 78.65 = ATCC 33071* | yes |
| *Rahnella sp. Y9602* | yes |
| *Rheinheimera nanhaiensis E407-8* | no |
| *Rheinheimera sp. A13L* | no |

| Genome | has F7 |
|---|---|
| *Rhodanobacter fulvus Jip2* | yes |
| *Rhodanobacter sp. 116-2* | no |
| *Rhodanobacter spathiphylli B39* | no |
| *Rhodanobacter thiooxydans LCS2* | yes |
| *Saccharophagus degradans 2-40* | yes |
| *Salinisphaera shabanensis E1L3A* | yes |
| *Salmonella bongori N268-08* | yes |
| *Salmonella enterica subsp. enterica serovar Gallinarum str. 9184* | yes |
| *Serratia liquefaciens ATCC 27592* | yes |
| *Serratia marcescens VGH107* | yes |
| *Serratia odorifera 4Rx13* | yes |
| *Serratia plymuthica S13* | yes |
| *Serratia proteamaculans 568* | yes |
| *Serratia sp. AS13* | yes |
| *Shewanella amazonensis SB2B* | yes |
| *Shewanella baltica OS155* | yes |
| *Shewanella benthica KT99* | yes |
| *Shewanella denitrificans OS217* | no |
| *Shewanella frigidimarina NCIMB 400* | no |
| *Shewanella halifaxensis HAW-EB4* | no |
| *Shewanella loihica PV-4* | yes |
| *Shewanella oneidensis MR-1* | yes |
| *Shewanella pealeana ATCC 700345* | no |
| *Shewanella piezotolerans WP3* | no |
| *Shewanella putrefaciens CN-32* | no |
| *Shewanella sediminis HAW-EB3* | yes |

| **Genome** | **has F7** |
| --- | --- |
| *Shewanella sp. MR-4* | yes |
| *Shewanella violacea DSS12* | yes |
| *Shewanella woodyi ATCC 51908* | yes |
| *Shigella boydii CDC 3083-94* | yes |
| *Shigella dysenteriae 1617* | no |
| *Shigella flexneri 4343-70* | yes |
| *Shigella sonnei Ss046* | yes |
| *Shigella sp. D9* | yes |
| *Simiduia agarivorans SA1 = DSM 21679* | yes |
| *Stenotrophomonas maltophilia K279a* | yes |
| *Stenotrophomonas sp. SKA14* | yes |
| *Teredinibacter turnerae T7901* | yes |
| *Thalassolituus oleivorans MIL-1* | yes |
| *Thioalkalimicrobium aerophilum AL3* | yes |
| *Thioalkalivibrio sp. K90mix* | no |
| *Thioalkalivibrio sulfidiphilus HL-EbGr7* | no |
| *Thiocapsa marina 5811* | no |
| *Thiocystis violascens DSM 198* | yes |
| *Thiomicrospira crunogena XCL-2* | yes |
| *Thiorhodococcus drewsii AZ1* | yes |
| *Thiorhodospira sibirica ATCC 700588* | yes |
| *Thiorhodovibrio sp. 970* | no |
| *Thiothrix nivea DSM 5205* | no |
| *Vibrio alginolyticus 40B* | no |
| *Vibrio anguillarum 775* | yes |
| *Vibrio brasiliensis LMG 20546* | yes |

| Genome | has F7 |
|---|---|
| *Vibrio campbellii CAIM 519 = NBRC 15631* | no |
| *Vibrio caribbenthicus ATCC BAA-2122* | no |
| *Vibrio cholerae HC-23A1* | no |
| *Vibrio coralliilyticus ATCC BAA-450* | yes |
| *Vibrio fischeri MJ11* | no |
| *Vibrio furnissii CIP 102972* | yes |
| *Vibrio harveyi 1DA3* | no |
| *Vibrio ichthyoenteri ATCC 700023* | no |
| *Vibrio metschnikovii CIP 69.14* | no |
| *Vibrio mimicus MB451* | yes |
| *Vibrio nigripulchritudo ATCC 27043* | yes |
| *Vibrio ordalii ATCC 33509* | yes |
| *Vibrio orientalis CIP 102891 = ATCC 33934* | yes |
| *Vibrio parahaemolyticus O1:Kuk str. FDA_R31* | no |
| *Vibrio rotiferianus DAT722* | no |
| *Vibrio scophthalmi LMG 19158* | no |
| *Vibrio shilonii AK1* | no |
| *Vibrio sinaloensis DSM 21326* | yes |
| *Vibrio sp. HENC-01* | no |
| *Vibrio splendidus ATCC 33789* | no |
| *Vibrio tubiashii NCIMB 1337 = ATCC 19106* | yes |
| *Vibrio vulnificus MO6-24/O* | yes |
| *Vibrionales bacterium SWAT-3* | no |
| *Wohlfahrtiimonas chitiniclastica SH04* | no |
| *Xanthomonas albilineans GPE PC73* | yes |
| *Xanthomonas axonopodis pv. malvacearum str. GSPB2388* | yes |

| Genome | has F7 |
|---|---|
| *Xanthomonas campestris pv. musacearum NCPPB 4381* | yes |
| *Xanthomonas citri subsp. citri Aw12879* | yes |
| *Xanthomonas fuscans subsp. aurantifolii str. ICPB 10535* | yes |
| *Xanthomonas gardneri ATCC 19865* | yes |
| *Xanthomonas oryzae pv. oryzicola BLS256* | yes |
| *Xanthomonas perforans 91-118* | yes |
| *Xanthomonas sacchari NCPPB 4393* | yes |
| *Xanthomonas translucens DAR61454* | yes |
| *Xanthomonas vesicatoria ATCC 35937* | yes |
| *Xenorhabdus bovienii SS-2004* | yes |
| *Xenorhabdus nematophila ATCC 19061* | yes |
| *Xylella fastidiosa Temecula1* | no |
| *Yersinia aldovae ATCC 35236* | yes |
| *Yersinia bercovieri ATCC 43970* | yes |
| *Yersinia enterocolitica subsp. palearctica Y11* | yes |
| *Yersinia frederiksenii ATCC 33641* | yes |
| *Yersinia intermedia ATCC 29909* | yes |
| *Yersinia kristensenii ATCC 33638* | yes |
| *Yersinia mollaretii ATCC 43969* | yes |
| *Yersinia pestis PY-16* | yes |
| *Yersinia pseudotuberculosis PB1/+* | yes |
| *Yersinia rohdei ATCC 43380* | yes |
| *Yersinia ruckeri ATCC 29473* | yes |
| *Yokenella regensburgei ATCC 43003* | yes |
| *endosymbiont of Riftia pachyptila (vent Ph05)* | yes |
| *gamma proteobacterium HdN1* | no |

**Table S9.**
Genomes used in phylogenetic profiles.

| **Genomes imaged in this study** |
|---|
| *Methylomicrobium alcaliphilum 20Z* |
| *Pseudomonas aeruginosa PAO1* |
| *Shewanella oneidensis MR-1* |
| *Vibrio cholerae O1 biovar El Tor str. N16961* |

| **Gamma-Proteobacteria** |
|---|
| *Acinetobacter baumannii AB0057* |
| *Acinetobacter calcoaceticus PHEA-2* |
| *Acinetobacter oleivorans DR1* |
| *Aeromonas hydrophila subsp. hydrophila ATCC 7966* |
| *Aeromonas salmonicida subsp. salmonicida A449* |
| *Aeromonas veronii B565* |
| *Alcanivorax borkumensis SK2* |
| *Alcanivorax dieselolei B5* |
| *Aliivibrio salmonicida LFI1238* |
| *Alkalilimnicola ehrlichii MLHE-1* |
| *Allochromatium vinosum DSM 180* |
| *Alteromonas macleodii str. 'Ionian Sea U7'* |
| *Alteromonas sp. SN2* |
| *Azotobacter vinelandii CA6* |
| *Cellvibrio japonicus Ueda107* |
| *Chromohalobacter salexigens DSM 3043* |
| *Citrobacter koseri ATCC BAA-895* |
| *Citrobacter rodentium ICC168* |
| *Colwellia psychrerythraea 34H* |
| *Cronobacter sakazakii ATCC BAA-894* |

| **Gamma-Proteobacteria** |
|---|
| *Cronobacter turicensis z3032* |
| *Dichelobacter nodosus VCS1703A* |
| *Dickeya dadantii Ech703* |
| *Dickeya zeae Ech1591* |
| *Edwardsiella ictaluri 93-146* |
| *Edwardsiella tarda C07-087* |
| *Enterobacter aerogenes KCTC 2190* |
| *Enterobacter asburiae LF7a* |
| *Enterobacter cloacae subsp. cloacae NCTC 9394* |
| *Enterobacter sp. 638* |
| *Enterobacteriaceae bacterium strain FGI 57* |
| *Erwinia amylovora ATCC 49946* |
| *Erwinia billingiae Eb661* |
| *Erwinia pyrifoliae Ep1/96* |
| *Erwinia sp. Ejp617* |
| *Erwinia tasmaniensis Et1/99* |
| *Escherichia coli O157:H7 str. EDL933* |
| *Escherichia fergusonii ATCC 35469* |
| *Ferrimonas balearica DSM 9799* |
| *Frateuria aurantia DSM 6220* |
| *Gammaproteobacteria gamma proteobacterium HdN* |
| *Glaciecola nitratireducens FR1064* |
| *Glaciecola psychrophila 170* |
| *Glaciecola sp. 4H-3-7+YE-5* |
| *Hahella chejuensis KCTC 2396* |
| *Halomonas elongata DSM 2581* |

| **Gamma-Proteobacteria** |
|---|
| *Halorhodospira halophila SL1* |
| *Halothiobacillus neapolitanus c2* |
| *Herminiimonas arsenicoxydans* |
| *Idiomarina loihiensis GSL 199* |
| *Kangiella koreensis DSM 16069* |
| *Legionella longbeachae NSW150* |
| *Listonella anguillarum M3* |
| *Marinobacter adhaerens HP15* |
| *Marinobacter aquaeolei VT8* |
| *Marinobacter hydrocarbonoclasticus ATCC 49840* |
| *Marinobacter sp. BSs20148* |
| *Marinomonas mediterranea MMB-1* |
| *Marinomonas posidonica IVIA-Po-181* |
| *Marinomonas sp. MWYL1* |
| *Methylococcus capsulatus str. Bath* |
| *Methylomonas methanica MC09* |
| *Methylophaga sp. JAM1* |
| *Morganella morganii subsp. morganii KT* |
| *Nitrosococcus halophilus Nc4* |
| *Nitrosococcus oceani ATCC 19707* |
| *Nitrosococcus watsonii C-113* |
| *Oceanimonas sp. GK1* |
| *Pantoea ananatis LMG 20103* |
| *Pantoea sp. At-9b* |
| *Pantoea vagans C9-1* |
| *Pectobacterium atrosepticum SCRI1043* |

| **Gamma-Proteobacteria** |
|---|
| *Pectobacterium carotovorum subsp. carotovorum PCC21* |
| *Pectobacterium sp. SCC3193* |
| *Pectobacterium wasabiae WPP163* |
| *Photobacterium profundum SS9* |
| *Photorhabdus asymbiotica* |
| *Photorhabdus luminescens subsp. laumondii TTO1* |
| *Proteus mirabilis BB2000* |
| *Providencia stuartii MRSN 2154* |
| *Pseudoalteromonas atlantica T6c* |
| *Pseudoalteromonas haloplanktis TAC125* |
| *Pseudoalteromonas sp. SM9913* |
| *Pseudomonas aeruginosa PA1* |
| *Pseudomonas brassicacearum subsp. brassicacearum NFM421* |
| *Pseudomonas denitrificans ATCC 13867* |
| *Pseudomonas entomophila L48* |
| *Pseudomonas fluorescens A506* |
| *Pseudomonas fulva 12-X* |
| *Pseudomonas mendocina ymp* |
| *Pseudomonas monteilii SB3101* |
| *Pseudomonas poae RE*1-1-14* |
| *Pseudomonas protegens Pf-5* |
| *Pseudomonas putida BIRD-1* |
| *Pseudomonas resinovorans NBRC 106553* |
| *Pseudomonas sp. TKP* |
| *Pseudomonas stutzeri DSM 4166* |
| *Pseudomonas syringae pv. syringae B728a* |

| **Gamma-Proteobacteria** |
|---|
| *Pseudoxanthomonas spadix BD-a59* |
| *Pseudoxanthomonas suwonensis 11-1* |
| *Psychrobacter arcticus 273-4* |
| *Psychrobacter cryohalolentis K5* |
| *Psychrobacter sp. PRwf-1* |
| *Psychromonas sp. CNPT3* |
| *Rahnella aquatilis CIP 78.65 = ATCC 33071* |
| *Rahnella sp. Y9602* |
| *Rhodanobacter sp. 2APBS1* |
| *Saccharophagus degradans 2-40* |
| *Salmonella bongori NCTC 12419* |
| *Salmonella enterica subsp. enterica serovar Typhi str. Ty21a* |
| *Serratia liquefaciens ATCC 27592* |
| *Serratia marcescens WW4* |
| *Serratia plymuthica AS9* |
| *Serratia proteamaculans 568* |
| *Serratia sp. AS12* |
| *Shewanella amazonensis SB2B* |
| *Shewanella baltica BA175* |
| *Shewanella denitrificans OS217* |
| *Shewanella frigidimarina NCIMB 400* |
| *Shewanella halifaxensis HAW-EB4* |
| *Shewanella loihica PV-4* |
| *Shewanella pealeana ATCC 700345* |
| *Shewanella piezotolerans WP3* |
| *Shewanella putrefaciens 200* |

| **Gamma-Proteobacteria** |
| --- |
| *Shewanella sediminis HAW-EB3* |
| *Shewanella sp. ANA-3* |
| *Shewanella violacea DSS12* |
| *Shewanella woodyi ATCC 51908* |
| *Shigella boydii CDC 3083-94* |
| *Shigella flexneri 2a str. 301* |
| *Shigella sonnei Ss046* |
| *Simiduia agarivorans SA1 = DSM 21679* |
| *Stenotrophomonas maltophilia JV3* |
| *Teredinibacter turnerae T7901* |
| *Thalassolituus oleivorans MIL-1* |
| *Thioalkalivibrio sp. K90mix* |
| *Thioalkalivibrio sulfidophilus HL-EbGr7* |
| *Thiocystis violascens DSM 198* |
| *Thiomicrospira crunogena XCL-2* |
| *Vibrio alginolyticus NBRC 15630 = ATCC 17749* |
| *Vibrio anguillarum 775* |
| *Vibrio campbellii ATCC BAA-1116* |
| *Vibrio cholerae O395* |
| *Vibrio fischeri ES114* |
| *Vibrio furnissii NCTC 11218* |
| *Vibrio harveyi ATCC BAA-1116* |
| *Vibrio nigripulchritudo* |
| *Vibrio parahaemolyticus RIMD 2210633* |
| *Vibrio sp. EJY3* |
| *Vibrio splendidus LGP32* |

| **Gamma-Proteobacteria** |
|---|
| *Vibrio vulnificus CMCP6* |
| *Xanthomonas albilineans GPE PC73* |
| *Xanthomonas axonopodis pv. citrumelo F1* |
| *Xanthomonas campestris pv. vesicatoria str. 85-10* |
| *Xanthomonas citri subsp. citri Aw12879* |
| *Xanthomonas oryzae pv. oryzae KACC 10331* |
| *Xenorhabdus bovienii SS-2004* |
| *Xenorhabdus nematophila ATCC 19061* |
| *Xylella fastidiosa subsp. fastidiosa GB514* |
| *Yersinia enterocolitica subsp. enterocolitica 8081* |
| *Yersinia pestis Antiqua* |
| *Yersinia pseudotuberculosis YPIII* |

| **Beta-Proteobacteria** |
|---|
| *Achromobacter xylosoxidans NBRC 15126 = ATCC 27061* |
| *Acidithiobacillus caldus SM-1* |
| *Bordetella pertussis 18323* |
| *Candidatus Accumulibacter phosphatis clade IIA str. UW-1* |
| *Collimonas fungivorans Ter331* |
| *Gallionella capsiferriformans ES-2* |
| *Janthinobacterium sp. Marseille* |
| *Ralstonia pickettii 12J* |
| *Ralstonia solanacearum Po82* |
| *Variovorax paradoxus S110* |

**Data S1. (separate file)**
Files used to produce the homology models as described in Table S7.

**Data S2. (separate file)**
Phylogenetic trees in Figure 4, S1, S2 and S4.
